# Connectome-Based Predictive Modeling of Concurrent and Prospective Substance Use in Adolescence

**DOI:** 10.1101/2025.09.01.673428

**Authors:** João F. Guassi Moreira, Nicholas Allgaier, Micah E. Johnson, Alexandra Potter, Hugh Garavan, Damien Fair

**Author notes:** Corresponding Author Brogden Psychology Building 1202 W Johnson St. Madison, WI 53706.

## Abstract

Understanding the neural mechanisms of adolescent substance use is a critical public health issue, with direct implications for bolstering prevention and treatment strategies. Yet this effort is challenging because substance use is multi-faceted, commonly used brain network features are not optimized to capture both local and global aspects of intrinsic connectivity, and because substance use facets change over time. In this study, we operationalized adolescent substance use along three dimensions—intent, access, and family-developmental history—and trained predictive models of each facet using resting-state connectivity. Trait impulsivity, a known risk factor, was also examined.

Using Baseline and 2 Year Follow-Up data from the ABCD Bids Community Collection (ABCC), we found that prediction was more successful at follow-up than baseline. At baseline, predictive accuracy was modest and intent to use substances was the most accurately predicted facet. Prediction accuracies at follow-up were much higher, with access and family-developmental history being better predicted, signaling a developmental shift in the brain–behavior mapping of substance use vulnerabilities.

These findings suggest that the neurobiological correlates of substance use are dynamic across adolescence, possibly reflecting changing phenotype. More broadly, these results underscore the importance of modeling distinct substance use facets and accounting for developmental timing to understand risk trajectories, while contributing to a growing literature that shows early-developing individual differences are predictive of later outcomes.

## 1. Introduction

The societal costs of substance use are enormous. In the United States alone, the economic burden of alcohol, tobacco, and illicit substances stretches upwards of $700 billion (Rehm et al., 2009; NIDA, 2017). Consequently, a central goal in translational neuroscience is to identify neurobiological markers that can predict substance use and related vulnerabilities. This problem is highly complex. Researchers must (i) determine how to operationalize substance use, (ii) grapple with technical challenges related to selecting specific facets of brain activity to measure and which modeling framework to pair them with, and (iii) build predictive models that account for developmental context in relevant phenotypes. In the current study, we address these challenges by defining substance use as a multidimensional construct, representing intrinsic brain networks using connectome embeddings that capture local and global topology, and evaluating predictive models across multiple developmental timepoints to test their sensitivity to age-related changes in substance use phenotypes.

### 1.1 Background

#### 1.1.1 Operationalizing Substance Use

The etiology of substance use is myriad, with available literature showing that various factors play a role in shaping use. Foundational genetic variation such as polymorphisms in dopamine-related genes ((Mallard et al., 2016; Ruchkin et al., 2021; Skowronek et al., 2006) have been linked to increased risk. Contextual-environmental factors such as family climate, socioeconomic status, and adverse early experiences appear to dynamically shape substance use tendencies throughout the lifespan (Dick et al., 2007; McGue et al., 2000; Rogers et al., 2022; Silberg et al., 2003). Similarly, sociocultural customs related to social group norms (Fujimoto & Valente, 2012; Hussong, 2002; Lilja et al., 2003) and substance availability (Steen, 2010; Toumbourou et al., 2007) represent additional vulnerabilities to substance use. Notably, these influences on substance use operate through equifinal pathways (Cicchetti & Rogosch, 1996) such that the same behavioral outcomes can be traced back onto diverse developmental histories, suggesting that studying substance use phenotypes and associated vulnerabilities in the collective may be a particularly fruitful method of inquiry.

For these reasons, defining substance use phenotypes in the quest for neurobiological markers is challenging. Even researchers who are interested in use of a single substance, or a particular substance use disorder (e.g., cannabis use disorder), must grapple with the complexity of isolating relevant behavioral phenotypes (such as frequency of use, dosage, delivery method, etc.) and account for the fact that use behaviors and vulnerabilities are often co-occurring across substances (Tomczyk et al., 2016; Vergunst et al., 2022), or that individuals may use substances in such a manner that does not meet clinical diagnostic thresholds but may nevertheless still carry important developmental, health, and societal consequences. Moreover, vulnerability to substance use is oft characterized by presence of transdiagnostic risk factors that are not intrinsically defined in relationship to substance use, but nevertheless evince strong associations with use (e.g., trait impulsivity). Because these vulnerabilities are often of interest in prevention, screening, or treatment efforts, their inclusion is warranted in the search for neurobiological markers.

### 1.1.2 Technical Hurdles: Measuring and Modeling with Connectivity

Resting state functional magnetic resonance imaging (rs-fMRI) is a putatively fruitful tool for identifying and understanding neurobiological precursors given its ability to assess core, intrinsic brain networks that are thought to underlie clinically relevant behaviors, cognitions, and motivations (Canario et al., 2021; Petersen et al., 2024; Schettino et al., 2024; Seitzman et al., 2019). The past quarter century has seen a massive surge of interest specifically in using brain network connectivity metrics derived from rs-fMRI as a means for understanding brain-behavior relationships a la predictive modeling (Ooi et al., 2025; Shen et al., 2017; Spisak et al., 2023). For instance, in the context of substance use research, recent work has shown that longitudinal resting-state functional connectivity patterns measured in late childhood and early adolescence prospectively differentiate youth who initiate substance use from matched controls in later follow-ups, and that these connectivity patterns are associated with environmental exposures such as pollution (Kardan et al., 2025). For as much optimism as there is in leveraging rs-fMRI to model substance use, however there is an accompanying basket of technical hurdles (Uddin et al., 2024; Wu et al., 2023).

One hurdle lies in how functional brain networks are represented. Traditional approaches rely on pairwise Pearson correlations between the functional time series of brain regions. These metrics are conceptually straightforward but can be noisy and may fail to capture higher-order network structure (Bastos & Schoffelen, 2015; Cutts et al., 2023, 2025; Milisav et al., 2024). This is because brain networks contain both *local* information (connections amongst specific pairs of regions) and *global* topographical information (overall network architecture) (Petersen & Sporns, 2015; Rosenthal et al., 2018). Focusing more one type of information comes at the risk of discarding valuable information about the other. Newer available methods such as connectome embeddings—vector representations learned from network topology by fitting high dimensional computational models—aim to encode both local and global features in a parsimonious form (Rosenthal et al., 2018; Levakov et al., 2018), but have not yet been used extensively in connectome-based predictive modeling.

A second hurdle relates to modeling choices, specifically the use of predictive modeling (e.g., using cross-validation to build a model to predict new, unseen data) versus descriptive approaches (e.g., testing pairwise associations between brain data and phenotypes of interest). While historical attempts at identifying neurobiological markers have favored descriptive models due to their conceptual tractability and ostensible utility for generating causal insights (Yarkoni & Westfall, 2017), they tend to be statistically inefficient and more vulnerable to noise (Spisak et al., 2023).

Consequently, brain-behavior phenotypes derived from such approaches often generalize less effectively compared to those identified through predictive modeling methods (Marek et al., 2019, 2022; Spisak et al., 2023; Yarkoni & Westfall, 2017).

#### 1.1.3. Accounting for Developmental Context

Identifying neurobiological markers of substance use is a developmental science problem. Substance use trajectories often begin within the first two decades of life and indicators of substance use undergo meaningful change as well.

The onset of substance use and related vulnerabilities often emerge in adolescence. This phase in the lifespan is characterized, among other things, by the emergence of heightened risk-taking propensity (Duell et al., 2018; Tervo-Clemmens et al., 2024). While some kinds of risk-taking can potentially facilitate positive developmental outcomes (Do et al., 2017; Duell & Steinberg, 2019), others like substance use can seriously threaten adolescent wellbeing (Cunningham et al., 2018). Substance use in adolescence is increasingly prevalent—approximately 25 to 50% of teens report substance use experimentation during high school (Johnston et al., 2019)—and is highly consequential for usage habits later in life (Centers for Disease Control and Prevention, 2020; Garofoli, 2020). Moreover, vulnerabilities to substance use, such as peer deviance or reduced parental monitoring, also manifest during this time (for instance, as teens become susceptible to peer influences and are granted more autonomy; Blakemore & Mills, 2014; Steinberg & Morris, 2001).

Further compounding this challenge is the fact that substance use and its related vulnerabilities are prone to developmental changes across the lifespan, meaning that relevant indicators can be expressed differently in early adolescence compared to mid adolescence compared to young adulthood. That substance use and related vulnerabilities shift during adolescence makes it imperative for predictive models to contend with changing behavioral baselines and developmental trajectories: predicting a phenotype at one age is not equivalent to predicting it years later. Testing whether predictive models generalize across developmental stages is therefore crucial.

### 1.2 Current Study

To address the three challenges enumerated here, we leveraged data from the ABCD Study (Casey et al., 2018), the largest longitudinal neurodevelopmental cohort available.

To grapple with the challenge of operationalizing substance use amidst considerable heterogeneity across many phenotypes, we followed an existing strategy by computing composite measures from a large set of substance use-related variables in the ABCD dataset (Rapuano et al., 2020): (i) *intent* (intentions to use or active use), (ii) *accessibility* (direct or indirect availability of substances), and (iii) *family-developmental history* (parental or familial substance use, including developmental exposure). We also examined trait impulsivity given its role as a strong transdiagnostic risk factor for substance use.

The challenge of balancing local and global structure in networks defined with rs-fMRI data was addressed by deriving network features from functional connectome embeddings (Rosenthal et al., 2018; Levakov et al., 2018). Connectome embeddings are designed to capture both local and global information about network topology in a parsimonious and efficient manner by training a shallow autoencoder neural network (node2vec, Levakov et al., 2018) on a traditional connectivity matrix to produce vector embeddings for each node in the network. By being able to capture information from both global and local network characteristics, embedding-derived features have the potential to enhance predictive modeling outcomes, potentially outperforming traditional metrics. Given the relatively recent advent of connectome embedding models and our novel application of them to predictive modeling in this context, we compared their performance to other connectivity metrics (observed connectivity calculated using a correlation coefficient and correlations implied by pairwise node embedding similarities) in the Supplement (Section S1). Embeddings were employed in a multivariate connectome-based predictive modeling framework to relate functional connectivity metrics to the three substance use facets and trait impulsivity. In tapping connectome-based predictive modeling, we hope to quantify brain-behavior associations in a way that is generative, statistically generalizable, and more robust to noise.

Finally, to deal with the developmentally-relevant nature of substance use, we trained and evaluated connectome-based predictive models on Baseline rs-fMRI data to independently predict each substance use composite and impulsivity at both Baseline and the 2 Year Follow Up. This design allowed us to assess how well brain features at baseline predict concurrent phenotypes and how predictive performance changes when the same facets are measured two years later. We chose to use rs-fMRI connectivity data from Baseline because there is notable interest in understanding early substance use susceptibility (Kardan et al., 2025; Jackson et al., 2015; Azagba et al., 2015; Green et al., 2024), recent work suggests early neural phenotypes are more consequential for downstream outcomes (Xie et al., 2025), and because other complementary work demonstrates that longitudinal resting-state connectivity can prospectively distinguish substance use initiators from non-initiators (Kardan et al., 2025), underscoring the developmental trajectory of brain networks and the promise of predictive approaches.

Overall, by integrating multidimensional behavioral phenotypes, complementary brain connectivity representations, and a multi-wave prediction framework, we aim to provide a more complete account of how intrinsic brain networks relate to substance use and its vulnerabilities in adolescence.

## 2. Methods

### 2.1 Participants

Data for the current project were taken from the ongoing Adolescent Brain and Cognitive Development (ABCD) study (Casey et al., 2018). The ABCD study is a large, multi-site longitudinal study designed to comprehensively study psychological and neurobiological development across the second decade of life, with emphasis placed on understanding links to addiction, substance use, and other varied wellbeing outcomes. Here we give a brief overview of ABCD study details relevant to the current project. In-depth details of study design, recruitment, and full description of measures are available elsewhere (Casey et al., 2018; Garavan et al., 2018). The study recruited over 11,875 children between the ages of 9 to 11 from 21 research sites across the United States and tracked them for several years. A battery of measures was collected from participants, including the rs-fMRI and substance use data used here. Participants were screened based on the quality of resting state fMRI data. We only included from participants who had 10 minutes of usable resting state data following acceptable head motion thresholds (see *2.2 fMRI Data Acquisition & Preprocessing*). In total, our sample was comprised of 5,955 participants (see Table 1 for demographics).

**Table 1.** Study Demographics Split by ABCC Arm.

| Variable | Arm 1 | Arm 2 |
| --- | --- | --- |
| *N* | 2,929 | 3,026 |
| Sex (percent female) | 52.75% | 50.63% |
| Ethnicity & Race |  |  |
| Hispanic/Latinx | 18.35% | 18.46% |
| Alaska Native | 0.00% | 0.00% |
| Asian Indian | 0.44% | 0.30% |
| Black | 12.51% | 13.14% |
| Chinese | 0.55% | 0.36% |
| Filipino | 0.31% | 0.23% |
| Guamanian | 0.00% | 0.00% |
| Japanese | 0.03% | 0.03% |
| Korean | 0.14% | 0.20% |
| Native American | 0.51% | 0.36% |
| Native Hawaiian | 0.00% | 0.00% |
| Other Asian | 0.21% | 0.30% |
| Other Pacific Islander | 0.10% | 0.10% |
| Samoan | 0.10% | 0.00% |
| Vietnamese | 0.14% | 0.10% |
| White | 69.57% | 68.37% |
| Other Race | 2.70% | 3.28% |
| Multi-Racial | 11.86% | 12.08% |
| Don't Know | 0.44% | 0.63% |
| Refused | 0.27% | 0.46% |
| Socioeconomic Status |  |  |
| Income | 7.50 (2.44) | 7.46 (2.23) |
| Parental Education | 17.39 (2.38) | 17.34 (2.47) |

*Note.* 'Arm' refers to the matched group variable from the ABCC collection. Income refers to total combined family income in the prior 12 months. Response options ranged from 1 to 10, corresponding to 1-'Less than \$5,000', 2-'\$5,000 - \$11,999', 3-'\$12,000 - \$15,999', 4-'\$16,000 - \$24,999', 5-'\$25,000 - \$34,999', 6-'\$35,000 - \$49,999', 7-'\$50,000 - \$74,999', 8-'\$75,000 - \$99,999', 9-'\$100,000 - \$199,999', 10-'\$200,000 and greater'. Respondents answering "don't know" or "decline to answer" were not included in the table mean. Parental education ranged from 0 ('Never attended/Kindergarten only') to 21 ('doctoral degree'); 17 corresponds to 'Associate degree: Academic program' and 18 corresponds to 'Bachelor's degree (i.e., B.A.)'

### 2.2 fMRI Data Acquisition & Preprocessing

At baseline, participants completed four resting state scan runs (five minutes each) with eyes open to help ensure at least ten minutes of data below accepted head motion thresholds. All participants were scanned with harmonized protocols across study sites. Further details about fMRI scanning protocols and procedures for the ABCD study have been documented at length elsewhere (Casey et al., 2018). The resting-state fMRI data used here were preprocessed by the ABCD-BIDS Community Collection (ABCC) team (Feczko et al., 2021) with the following additional pre-processing steps conducted by the Developmental Cognition and Neuroimaging Lab at the University of Minnesota. fMRI data were first de-trended and de-meaned over time based on low head-motion volumes with FD values that did not exceed 0.3 mm. Confound regression was then performed to remove several nuisance signals. This meant regressing the resting state timeseries against the mean time series for white matter, cerebrospinal fluid, and global signal, all translational and rotational motion parameters (12 total). The resulting timeseries was then bandpass filtered between 0.008 and 0.09 Hz using a second-order Butterworth filter applied in the forward and backward directions. Data from frames with a FD value greater than 0.3 mm were replaced with interpolated data from remaining frames to avoid re-introducing head motion artifacts (interpolated data discarded from analyses). The human connectome project (HCP) workbench software was used to convert CIFTI dense timeseries into parcellated timeseries following the 333 ROI Gordon parcellation (Gordon et al., 2016). An additional 19 subcortical ROIs were added for a total of 352 parcels. All greyordinate timeseries were then averaged within parcel. Functional connectivity matrices were calculated by taking the Pearson product moment correlation of all pairwise parcel timeseries.

### 2.3 Node Embedding of Functional Connectivity Data

Node embedding of functional connectivity was conducted using the `cepy` python package (Levakov et al., 2021). The package uses the node2vec autoencoder algorithm to estimate a vectorized representation of brain nodes that maintains their topological organization. In doing so, the algorithm is able to efficiently balance the capture of both local (edge-level) and global (network-level) information. This represents a potential improvement in multivariate connectome-based predictive modelling, as current approaches that rely on edgewise features may be disproportionately weighing local information.

Embedding the individual nodes (parcels) in a brain network is accomplished by simulating many sequences of random walks across the network (i.e., a sequence of nodes is generated probabilistically according to the ties among all nodes), recording the sequence of nodes along the walk, moving a sliding window over the sequence of nodes to finally predict the central node from its surrounding context nodes. The algorithm is fit by using an autoencoder, a type of fully connected artificial neural network that takes an input, embeds the individual elements into a latent space that preserves a low dimensional representation of the input, and then reconstructs an output.

Connectome embeddings were computed independently for each individual participant, resulting in a unique vector for each parcel. The vector represents the position of that parcel in an *n*-dimensional latent space, where distances between vectors reflect the topological relationships among regions in the functional connectome. Here we specified 30 latent dimensions for our embeddings (see the Supplement, Section S1 for full specification details). As described in Section 2.5.1, the dimension values for all vectors for all parcels were used in our model training and testing procedure. These embeddings theoretically provide an advantage over observed connectivity because of their ability to balance local versus global (edge vs network) level information as well as their parsimonious feature sets (30 latent dimensions x 352 parcels = 10,560 unique features compared to 61,776 unique features with a traditional connectivity matrix).

### 2.4 Operationalizing Substance Use and Trait Impulsivity

#### 2.4.1 Substance Use

We took a theory-driven, multidimensional approach to quantifying different facets of substance use in the ABCD dataset. Using a large pool of available substance use related items, we created three composite variables meant to capture distinct facets of substance use at Baseline: *intent, access*, and *family-developmental history*. The selection of these facets was motivated by both theoretical and empirical considerations. Empirically, this work builds directly on prior data-driven efforts to organize the high-dimensional substance-use variable space in the ABCD dataset (Rapuano et al., 2020), which identified latent structure among a large set of substance-related items using baseline data. We intentionally retained and extended this framework to ensure continuity with prior work and to avoid *ad hoc* selection of outcomes from a vast and heterogeneous measurement battery.

Conceptually, these three facets capture complementary and developmentally relevant aspects of substance use vulnerability: intent reflects curiosity, desire, and early engagement with substances (particularly appropriate in early adolescence when use is infrequent); access indexes contextual and social conditions that lower the threshold for use; and family-developmental history captures intergenerational and early environmental risk that is well-established in the substance use literature. Although Rapuano et al. (2020) identified two higher-order factors, inspection of the reported correlation structure suggested their solution was under-factored such that variables related to contextual access and familial history should be clustered, motivating our decision to treat them as distinct facets rather than collapsing them into a single dimension.

Alternative approaches—such as modeling all substance-related variables independently or including broader constructs (e.g., peer influence or psychiatric comorbidity)—were considered but ultimately not pursued. Many individual substance use items are sparse in early adolescence, making independent predictive modeling statistically inefficient and difficult to interpret. Moreover, variables such as peer influence or psychopathology are more appropriately treated as upstream risk factors rather than as facets of substance use itself.

Composites were computed for the Baseline and 2 Year Follow-Up time points. Items for the composites were identified by a previous study using Baseline data (Rapuano et al., 2020). For composite scores at Baseline, we used the same exact pool of items in the aforementioned study. The 2 Year Follow-Up composites were using the subset of baseline variables that were still collected at that time, as certain baseline variables were excluded from future time points because they were deemed developmentally irrelevant and therefore not collected again. The full lists of variables for each composite at each timepoint are provided in the Supplement (Section S2). The composite for each facet is described in greater detail below. All composite scores were calculated by averaging the relevant variables for each facet.

The *intent* composite reflects prior or ongoing substance use, curiosity about using substances, and desire to use substances. At baseline, this included items from the Lifetime Use Inventory (Lisdahl & Price, 2012), which first asks youths about whether they have *heard* of a substance, and then queries if they have *tried* or *regularly use* each substance via a timeline follow-back assessment (Cervantes et al., 1994).

Additional items related to this facet (assessing alcohol, nicotine, and caffeine use) were also available at baseline and thus included. The items about whether youth had heard of each substance were excluded from the 2 Year Follow-Up calculation of this composite because those items were no longer collected.

The *access* composite captures contextual and social factors that facilitate or encourage substance use, such as direct access to substances, lax parental rules or monitoring, and the presence of peers who promote substance use. This composite was based on peer group deviance items referencing substance use, household rules regarding use, and parental risk attitudes toward use. The items for this composite were the same in Baseline and 2 Year Follow-Up calculations since the entire set was collected at both timepoints.

Finally, the *family-developmental history (FDHX)* refers to family history or parental developmental history of substance use, including in utero exposure. For Wave 2, family history items were excluded because these do not change over time; all other items were retained.

To reduce the influence of any outlying cases, scores on each composite at both timepoints were winsorized at 3.5 standard deviations.

#### 2.4.2 Trait Impulsivity

Trait impulsivity in the ABCD study was measured with an abbreviated youth version of the UPPS-P Impulsive Behavior Scale (Watts et al., 2020). The twenty-item scale taps five theoretically-informed dimensions of impulsivity (lack of perseverance, lack of premeditation, sensation seeking, negative urgency, and positive urgency) using a 1 (agree strongly) to 4 (disagree strongly) Likert scale. Participants’ responses to each statement were summed into a single score, with higher scores indicating more impulsivity. Scores were winsorized using the same criterion as the substance use variables.

### 2.5 Analysis Plan

#### 2.5.1 Connectome-Based Predictive Modeling of Substance Use and Trait Impulsivity

We leveraged machine learning and a variant of connectome-based predictive modeling (Shen et al., 2017) to engineer predictive models of substance use facets and trait impulsivity at Baseline and the 2 Year Follow-Up using Baseline rs-fMRI data. This process entails identifying the most relevant network features via pairwise associations with a given outcome variable and then using said features to train and validate a predictive model via machine learning techniques (e.g., cross-validation). Of note, the multivariate variant of connectome-based predictive modeling we employ here is better suited for our purposes given the dependency structure present in the connectivity metrics of interest here (Adkinson et al., 2024; Gao et al., 2019).

Overall, we ran 48 unique specifications of predictive model training and validation by varying the outcome being predicted (4), whether the outcomes were at Baseline or Year 2 Follow-Up (2), data splitting into discovery and validation samples (2), and the type of machine learning model used (3). Below we describe the details for each type of specification, followed by the model training and validation procedure.

##### 2.5.1.1 Outcomes

The three facets of substance use—intent, access, family-developmental history—in addition to trait impulsivity were specified as dependent variables in modeling.

##### 2.5.1.2 Outcome Timing

Measures of substance use and trait impulsivity were taken at Baseline and the 2 Year Follow-Up.

##### 2.5.1.3 Data Splitting

We took advantage of ABCC’s `matched_group` variable that sorts participants into two arms matched on various demographics including race, ethnicity, sex, and socioeconomic status (Arm 1 *n* = 2929, Arm 2 *n* = 3026; https://nda-abcd-collection-3165.readthedocs.io/latest/recommendations/#2-the-bids-participants-files-and-matched-groups). The model training and validation process leveraged was repeated twice such that each arm was used as both a discovery dataset for feature selection and modeling training, and a confirmation dataset for validation (each across independent iterations).

We chose to use the pre-defined matched group designations from the ABCC collection for model training and validation because they afforded us demographically-matched independent sets. This is in contrast to other recent approaches that perform feature selection and cross-validation in the same sample over many repetitions (Adkinson et al., 2024). Without detracting from other such work, our approach theoretically affords us a relatively better estimate of out-sample predictive ability because the model training and validation sets are completely independent and thus minimize any possibility of test-train leakage.

##### 2.5.1.4 Model Type

We used three types of models: partial least squares regression (PLSR), ridge regression (RIDGE), and XGBoost (XGB). The former two were implemented in `sklearn` python package, whereas XGB was implemented with the `xgboost` python package (`XGBRegressor()` function). The PLSR and RIDGE models each contained one hyperparameter, the number of components (1 to 10) and λ (l2 penalty term; 1e-6 to 1e6 in 20 linearly spaced intervals). The XGB model specified four hyperparameters: the number of estimators (100 or 200), learning rate (0.01, 0.05, 0.10), max depth (3 or 5), and λ (1, 10, 100). Hyperparameters were rounded to whole integers when necessary (e.g., number of components, max depth).

##### 2.5.1.5 Procedure

For each outcome variable of substance use, the following procedure was enacted (all implemented with `sklearn` functions). First, the discovery sample was used for feature selection by chaining the `RobustScaler()` function with its `SelectKBest()`function into a pipeline that normalized the predictors and then identified the top 1,000 features evincing the strongest association with the given dependent variable via univariate regression. Next, once all features were selected, nested K-fold cross-validation (K = 5 outer folds) was used to train predictive models for each dependent variable in the discovery dataset. On each iteration of the inner loop, the `GridSearchCV()` function was used to optimize model hyperparameters by minimizing the mean squared error. The best hyperparameters from the grid search on each inner fold were saved as part of the outer loop partition. Once the procedure had been repeated for all outer folds, the best hyperparameters across all outer folds were averaged and used to estimate the model to the full discovery dataset. The model was then fit to the confirmation dataset and an *R^2^* value was created by correlating model predictions and observed results (Pearson’s *r*) and squaring the result. Models of the facets at the 2 Year Follow-Up controlled for baseline values of the modeled facet.

While we previously justified our choice to use connectome embeddings in our modeling procedure (e.g., capture of both edge-level and network-level connectivity characteristics while offering greater parsimony than full observed connectivity matrices) rather than traditional connectivity matrices, we also wanted to avoid the possibility that this choice inadvertently reduced model performance. To this end, we compared node embedding performance with traditional connectivity (i.e., observed connectivity), model-implied connectivity (i.e., pairwise cosine similarity among all encoding vectors), and two combined feature sets; these results are provided in the Supplement (Supplementary Tables 2-3).

#### 2.5.2 Non-fMRI Covariates

Non-fMRI covariates were added to each model as part of the feature selection process described above. An initial set of variables comprising data on biological sex, race, ethnicity and socioeconomic status was assembled from ABCD’s study demographic information. Biological sex (male, female) and ethnicity (Hispanic/Latinx vs not Hispanic/Latinx) were binary coded. Race was quantified as a set of one-hot encoded vectors for all the racial categories assessed in the ABCD study (Alaskan Native, Asian, Black, Chinese, Filipino, Guamanian, Hawaiian Native, Indian, Japanese, Korean, Native American, Samoan, Vietnamese, White, Other Asian, Other Pacific Islander, Other Race). This set also included variables for the ‘don’t know’ and ‘declined’ categories, in addition to a composite variable indicated mixed race status (defined as selecting more than one of the racial categories). Socioeconomic status was included because of its strong relationship to wellbeing across development (Peverill et al., 2021), and was captured by a set of variables comprised of parental education and household income.

The entire set of 23 demographic variables described above was concatenated with the discovery data for each iteration of the aforementioned modeling procedure and feature selection was performed as previously described over the entire set of fMRI and non-fMRI features. This procedure gave us the advantage of adjusting for potential non-brain confounds in a data-driven manner while potentially freeing up additional model degrees of freedom to be used on fMRI predictors. In addition to these demographic covariates, we also included site and scanner manufacturer in the same manner as described above.

## 3. Results

### 3.1 Establishing Fidelity of Connectome Embeddings

Our connectome embeddings faithfully represented functional connectivity as defined by the correlation of rs-fMRI parcel timeseries. Details about this analysis can be accessed in the Supplement (Section S1, Supplementary Figure 1).

### 3.2 Connectome-Based Prediction of Substance Use and Impulsivity

Full results of our connectome-based approach to modeling different facets of substance use and impulsivity at *Baseline* using five different feature sets are listed in Table 2. In line with the prior literature, we observed modest effect sizes that fell between the range of *r* = [0.001, 0.104] (*R^2^* = [0.00%,1.09%]), with most of the values typically falling between *r* = (0.05, 0.1). This means that even the best predictive models accounted for at most approximately 1% of the variance in substance use facets.

**Table 2.** Model performances predicting substance use composites at Baseline

| ***Intent*** |  |  |
| --- | --- | --- |
|  | *Arm 1* | *Arm 2* |
| PLSR | 0.095 (0.90%) | 0.090 (0.81%) |
| RIDGE | 0.069 (0.47%) | 0.104 (1.09%) |
| XGB | 0.076 (0.58%) | 0.086 (0.73%) |
| ***Access*** |  |  |
|  | *Arm 1* | *Arm 2* |
| PLSR | 0.075 (0.57%) | 0.056 (0.32%) |
| RIDGE | 0.059 (0.35%) | 0.071 (0.51%) |
| XGB | 0.072 (0.52%) | 0.072 (0.51%) |
| ***Family-Developmental History*** |  |  |
|  | *Arm 1* | *Arm 2* |
| PLSR | 0.059 (0.35%) | 0.091 (0.83%) |
| RIDGE | 0.085 (0.72%) | 0.085 (0.73%) |
| XGB | 0.089 (0.79%) | 0.093 (0.86%) |
| ***Impulsivity (UPPS)*** |  |  |
|  | *Arm 1* | *Arm 2* |
| PLSR | 0.035 (0.12%) | 0.039 (0.15%) |
| RIDGE | 0.041 (0.17%) | 0.050 (0.25%) |
| XGB | 0.029 (0.08%) | 0.017 (0.03%) |

*Note.* ‘Arm’ refers to the arm used for confirmation. ‘XGB’, ‘RIDGE’, and ‘PLSR’ refer to models trained and tested with XGBoost, ridge regression, and partial least squares regression respectively. Entries reflect Pearson correlation’s between fitted and observed values during model testing; parenteticals reflect the square of such values ( $R^2$ ) on a percent scale, indicating the proportion of variance explained. These results reflect features comprised of node embedding dimensions; comparisons against other feature sets can be accessed in the supplement.

Results for models predicting *2 Year Follow-Up* substance use and impulsivity are listed in Table 3. Consistent with recent work (Xie et al., 2025), predictive accuracy was generally higher when forecasting later outcomes from baseline rs-fMRI data, particularly for substance use composites. For intent, access, and family-developmental history, many models exceeded r = 0.10, with some—especially for family-developmental history—reaching values between 0.312 (*R^2^* = 9.71%) and 0.546 (*R^2^* = 29.82%). In contrast, prediction of Impulsivity remained weak, with all r values below 0.10.

**Table 3.** Model performances predicting substance use composites at the 2 Year Follow-Up.

| ***Intent*** |  |  |
| --- | --- | --- |
|  | *Arm 1* | *Arm 2* |
| PLSR | 0.110 (1.21%) | 0.104 (1.08%) |
| RIDGE | 0.100 (1.01%) | 0.116 (1.34%) |
| XGB | 0.180 (3.23%) | 0.193 (3.72%) |
| ***Access*** |  |  |
|  | *Arm 1* | *Arm 2* |
| PLSR | 0.268 (7.19%) | 0.177 (3.12%) |
| RIDGE | 0.156 (2.45%) | 0.143 (2.06%) |
| XGB | 0.394 (15.56%) | 0.393 (15.46%) |
| ***Family-Developmental History*** |  |  |
|  | *Arm 1* | *Arm 2* |
| PLSR | 0.312 (9.71%) | 0.378 (14.31%) |
| RIDGE | 0.446 (19.87%) | 0.516 (26.62%) |
| XGB | 0.504 (25.42%) | 0.546 (29.82%) |
| ***Impulsivity (UPPS)*** |  |  |
|  | *Arm 1* | *Arm 2* |
| PLSR | 0.048 (0.23%) | 0.082 (0.67%) |
| RIDGE | 0.027 (0.07%) | 0.031 (0.09%) |
| XGB | 0.031 (0.10%) | 0.092 (0.85%) |

*Note.* ‘Arm’ refers to the arm used for confirmation. ‘XGB’, ‘RIDGE’, and ‘PLSR’ refer to models trained and tested with XGBoost, ridge regression, and partial least squares regression respectively. Entries reflect Pearson correlation’s between fitted and observed values during model testing; parenteticals reflect the square of such values ( $R^2$ ) on a percent scale, indicating the proportion of variance explained. These results reflect features comprised of node embedding dimensions; comparisons against other feature sets can be accessed in the supplement.

We delve into our modeling results in more detail across the following subsections. Figures 1-3 depict modeling accuracy within each of the five modeling specifications; Figures 4-5 plot predicted versus observed scores for every individual modeling instance where matched group arm 1 was used as the confirmation sample (the same plots with arm 2 are included in the Supplement); Figures 6-7 depict the most important model features by whole-brain network (described in detail in Section 3.3).

**Figure 1.**
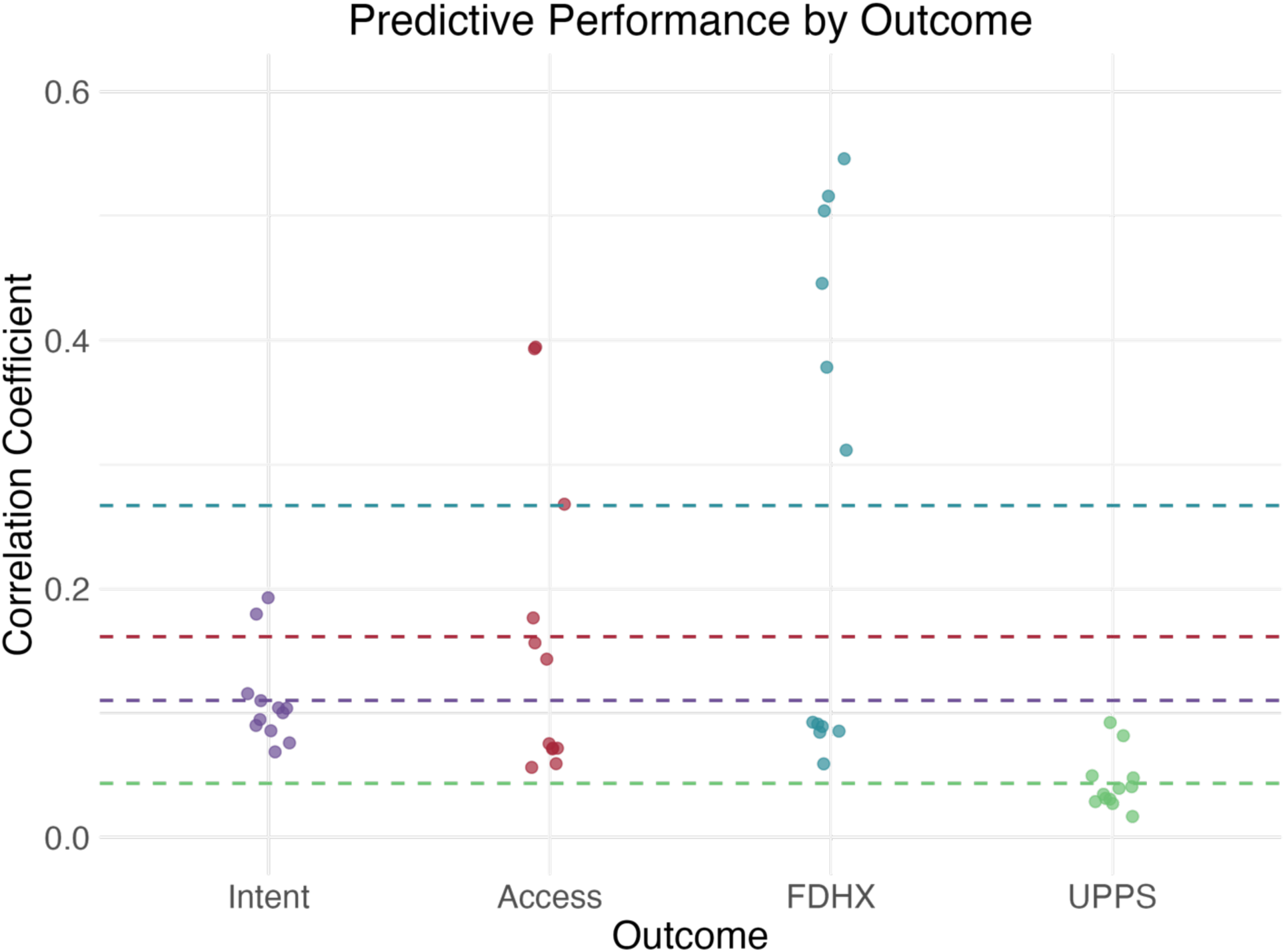
Predictive performances were highest for family-developmental history of substance use, collapsing across all other specifications

**Figure 2.**
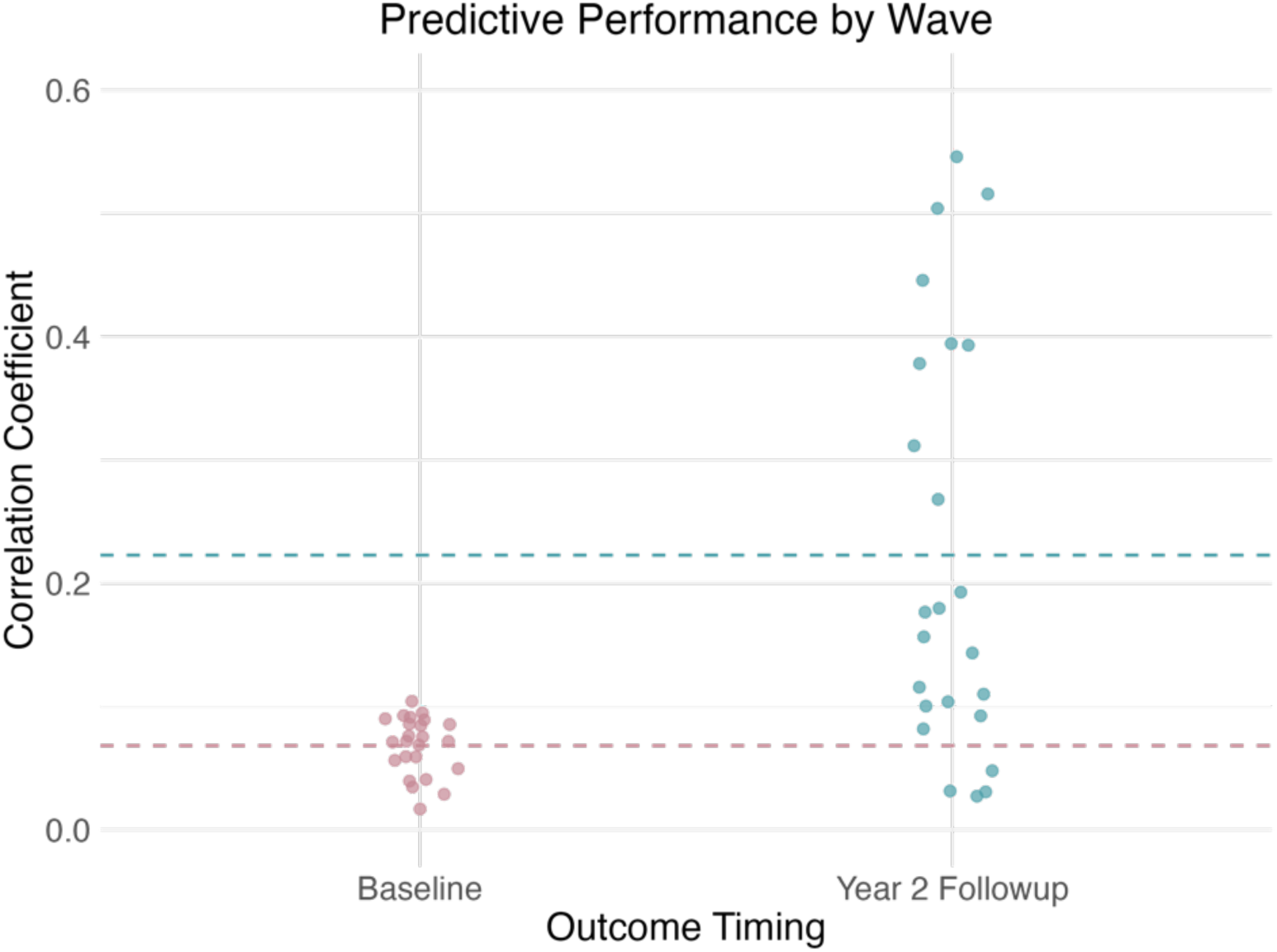
Predictive performances were highest for the 2 Year Follow-Up substance use, collapsing across all other specifications.

**Figure 3.**
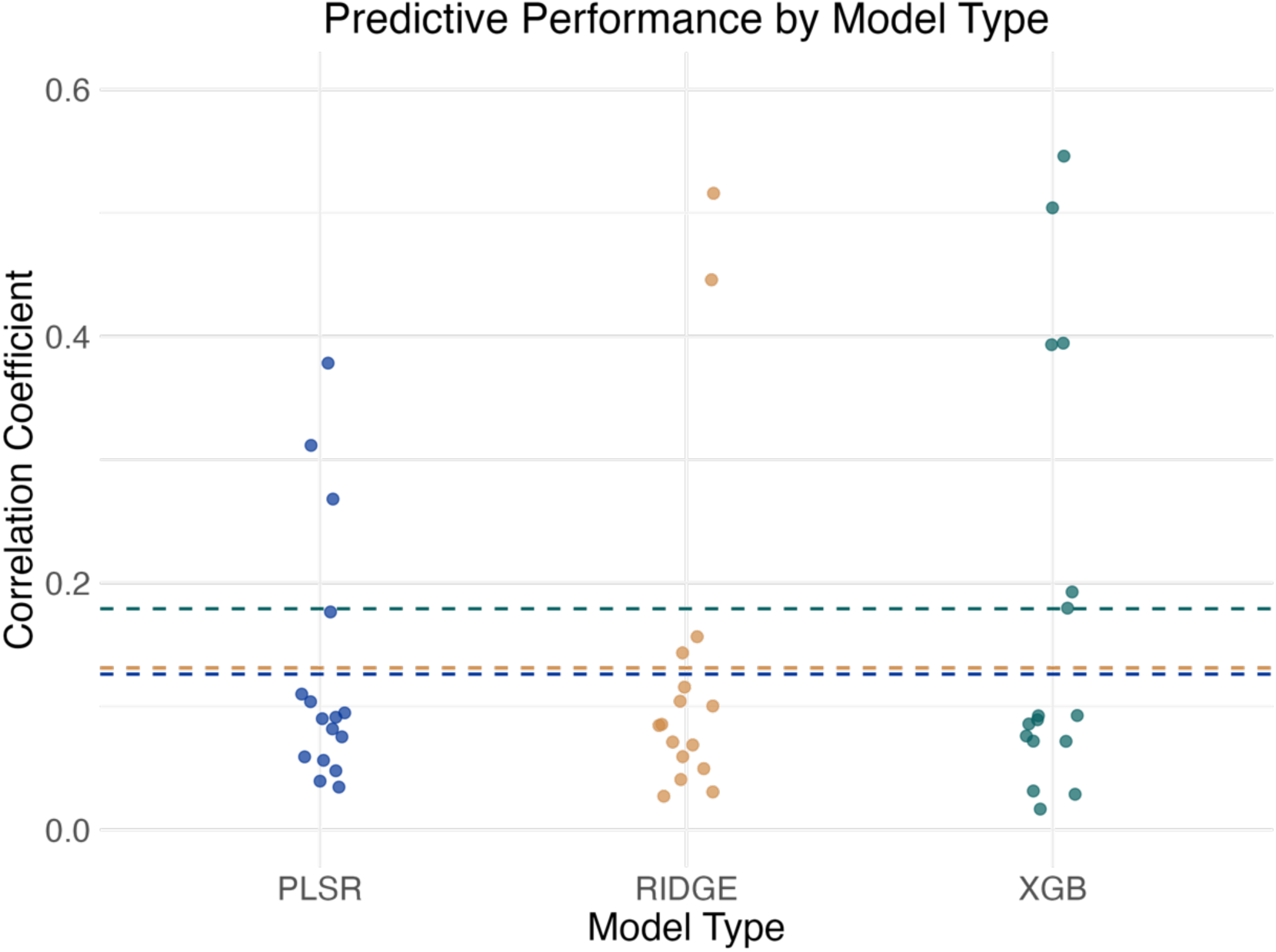
Predictive performances were comparable across the type of statistical model used, with a slight edge to XGBoost

**Figure 4.**
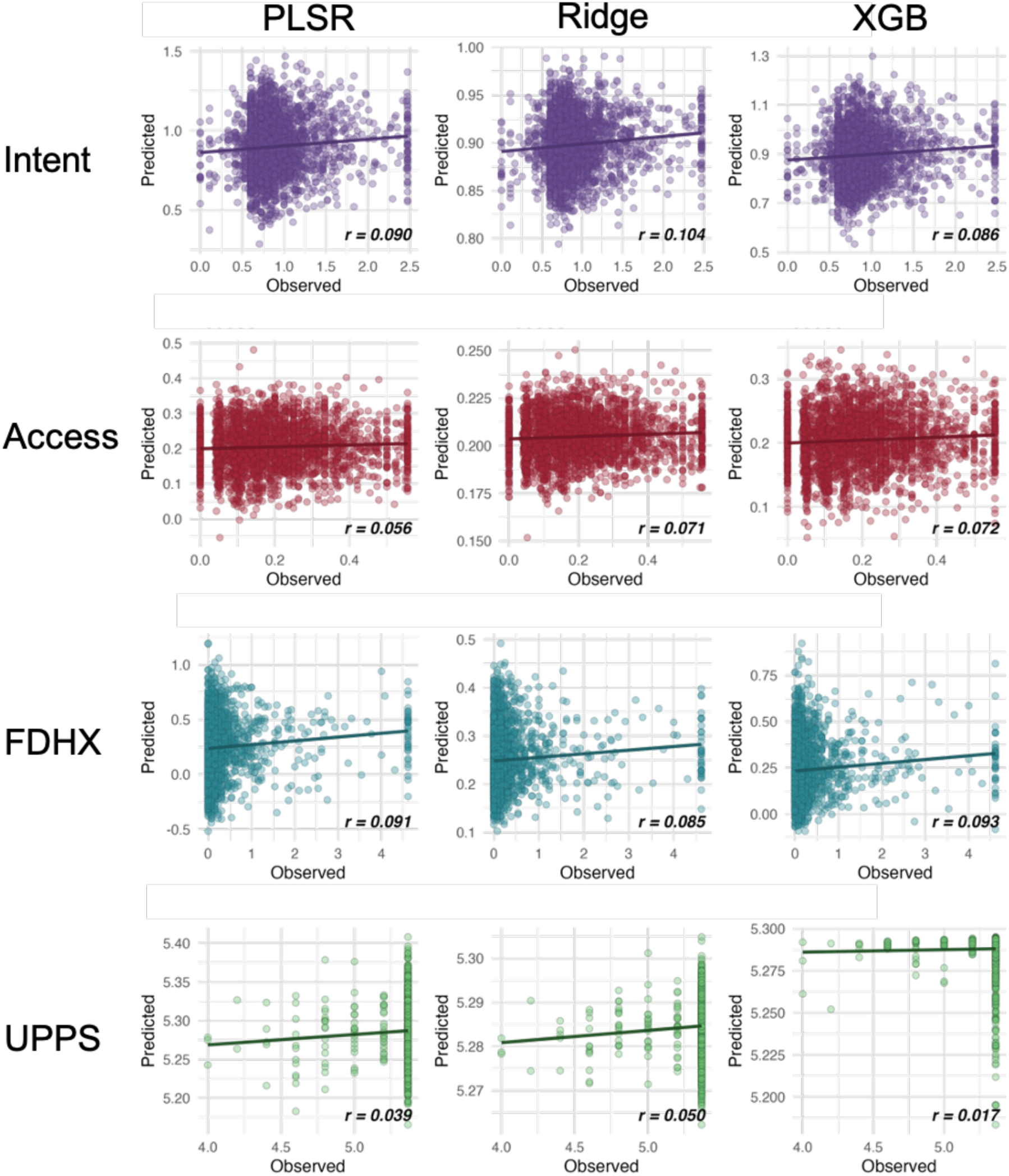
Model performance scatter plots by substance use facet at Baseline and model type.

**Figure 5.**
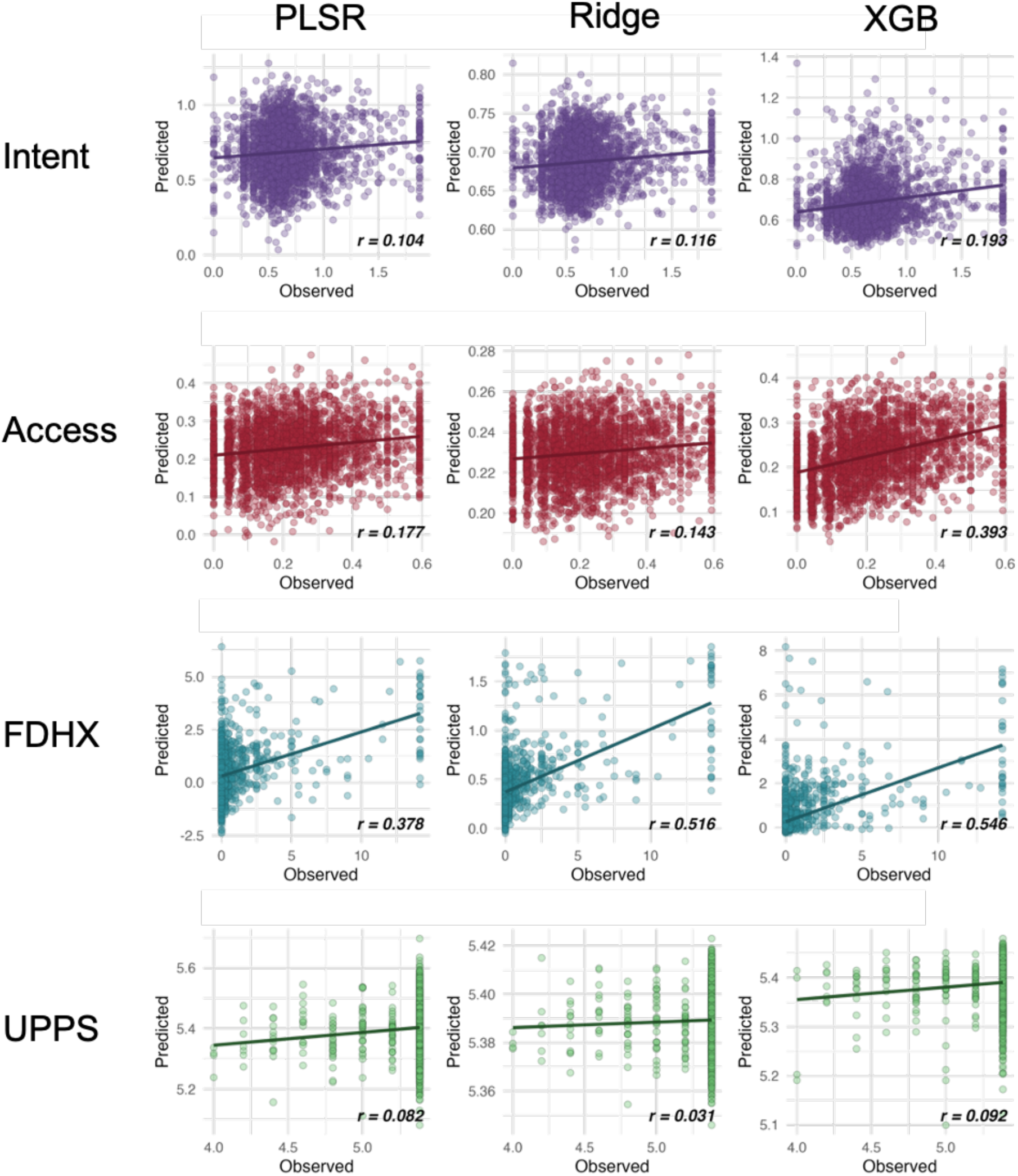
Model performance scatter plots by substance use facet at the 2 Year Follow-Up and model type.

**Figure 6.**
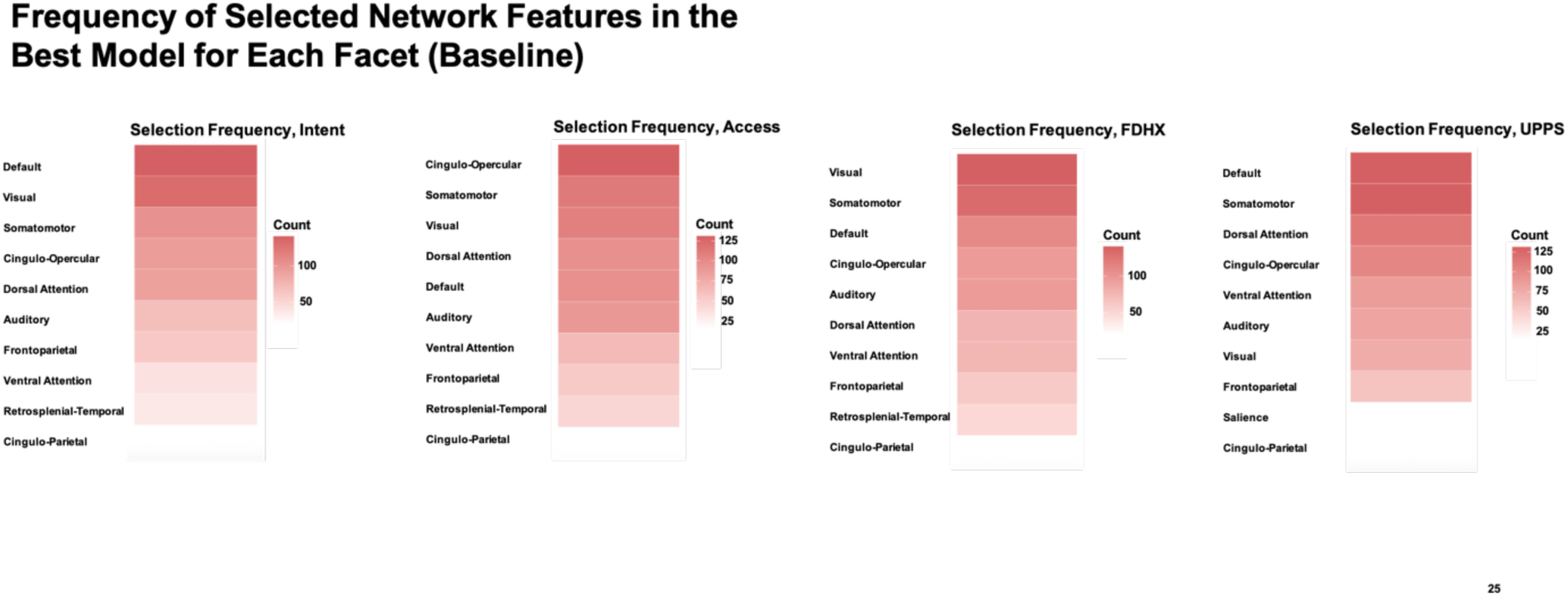
Most important networks, Baseline

**Figure 7.**
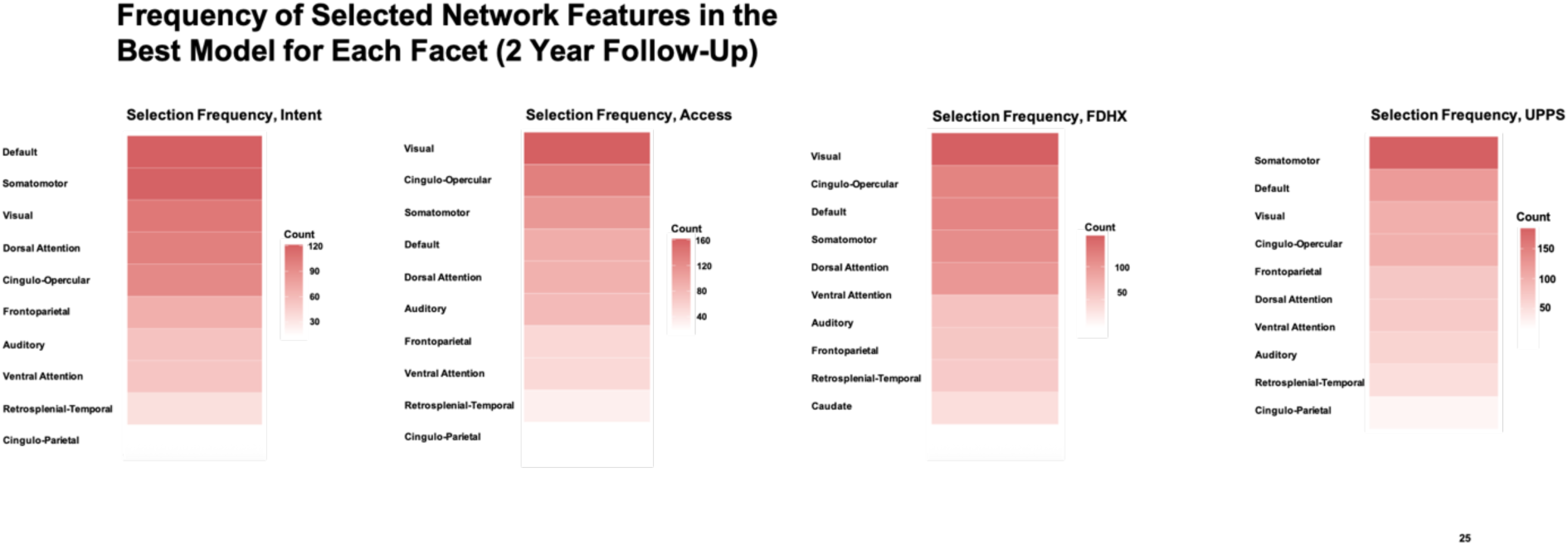
Most important networks, 2 Year Follow-Up

#### 3.2.1 Predictive Performance by Outcome

Model performance grouped by outcome (collapsing across all feature specifications and both timepoints) is depicted in Figure 1. Models were most accurate for family developmental history, followed by access, while prediction of impulsivity was consistently poor. These outcome-level differences align with patterns observed across both baseline and follow-up analyses (Tables 2 and 3). Notably, whereas baseline models performed best for intent, follow-up models showed substantially stronger prediction for access and family developmental history. Interpretations should be qualified by the fact that substance use composite definitions differed slightly between waves. In contrast to substance use facets, trait impulsivity was consistently poorly predicted at both baseline and follow-up, with all models yielding correlations below *r* = 0.10 (*R^2^* = 1%).

#### 3.2.2 Predictive Performance by Outcome Timing

Model performance grouped by outcome timing (collapsing across outcomes and feature specifications) is shown in Figure 2. Overall, predictive accuracy was markedly higher at the 2 Year Follow Up compared to baseline, with baseline models clustering near zero and follow-up models frequently exceeding *r* = 0.2 (*R^2^* = 4%). These differences were driven primarily by the substance use composites, as trait impulsivity was poorly predicted at both timepoints. Importantly, the outcome-specific results in Tables 2 and 3 clarify that this increase in accuracy was especially pronounced for family-developmental history and access, which showed strong improvements at follow-up, whereas Intent was better modeled at baseline.

#### 3.2.3 Predictive Performance by Machine Learning Model Type

Model performance grouped by machine learning model type (collapsing across outcomes, feature specifications, and timepoints) is shown in Figure 4. Predictive performance was broadly comparable between PLSR and ridge regression, both yielding modest accuracies. By contrast, XGBoost consistently outperformed the linear models, with average correlations higher by approximately 0.06.

#### 3.2.4 Supplementary Comparisons of Predictive Performance by Feature Set

Supplementary Figure 3 shows a validation analysis comparing node embeddings to alternative connectivity feature sets (observed connectivity, model-implied connectivity, and their combinations). These analyses were conducted to verify that node embeddings perform at least as well as traditional observed connectivity and model-implied connectivity, as well as combinations of these feature sets. Across all five specifications (node embeddings, observed connectivity, model-implied connectivity, observed connectivity + embeddings, and all features combined), predictive performance was broadly comparable. Importantly, feature sets that combined multiple connectivity metrics (e.g., all three metrics together, or observed connectivity plus embeddings) did not confer an accuracy advantage over node embeddings alone, indicating that node embeddings achieve predictive accuracy on par with traditional and combined feature sets, despite being a lower-dimensional representation of network topology at the data level.

#### 3.2.5 Predictive Performance by Sample Split

We checked to see whether model performance differed based on which matched group served as the confirmation arm. Supplementary Figure 5 shows predictive performance was comparable between sample splits.

#### 3.2.6 Post-Hoc Analyses

After obtaining our predictive modeling results, we ran two additional *post-hoc* analyses. First, we questioned whether low predictive performance for the Intent facet was driven by non-use variables (e.g., variables indicating curiosity or intent to use substances as opposed to items strictly about use). To address this, we trained predictive models to forecast an aggregate of use-only variables at the 2 Year Follow-Up (baseline use was not examined due to low prevalence). As shown in Supplementary Table 4, predictive performance for use alone was uniformly modest across feature sets, model types, and sample splits (*r*s generally < .07). Performance did not systematically exceed that observed for the intent composite at the same timepoint. These findings suggest that isolating substance use variables did not yield stronger predictive signal than modeling broader substance-related tendencies at this developmental stage.

Second, to assess whether increased predictive performance of the family-developmental history facet at the 2 Year Follow-Up was driven by different variable compositions at each timepoint (see Section 2.4.1), we re-computed the facet using an alternate calculation that incorporated baseline-only family history variables into the follow-up composite (see Supplementary Table 5). Predictive performance using this alternate composite was broadly comparable to the primary results, with some model specifications showing modest improvements, indicating that enhanced prediction at follow-up cannot be attributed solely to differences in composite variability.

### 3.3 Unpacking rs-fMRI Connectivity Features

Feature importance analyses revealed which large-scale brain networks contributed most strongly to model predictions (Figure 6 outcomes at Baseline; Figure 7 outcomes at the 2 Year Follow-Up). For these analyses, we examined the best-fitting model for each outcome served as the confirmation dataset (Arm 2 depicted for brevity). Embedding dimensions selected in the discovery split were mapped back to their corresponding parcels and further aggregated at the network level.

At Baseline, predictive features were relatively diffuse across networks. Given that baseline models generally showed weak predictive accuracy, this diffuse distribution likely reflects instability in feature selection rather than meaningful network-level contributions. By contrast, at the Year 2 Follow-Up, a clearer and more focused pattern emerged, with the Default Mode, Control, and Salience/Ventral Attention networks contributing disproportionately large numbers of features across the outcomes examined here.

## 4. Discussion

### 4.1 Overview

The present study sought to probe the underlying neurobiology of adolescent substance use by leveraging predictive modeling and resting state fMRI. In doing so, we aimed to address three key difficulties: (i) how to operationalize substance use, (ii) technical hurdles in rs-fMRI modeling—selecting and representing connectivity features and using predictive modeling rather than descriptive approaches—and (iii) capturing phenotypes that change with development. To that end, we partitioned substance-use measures into three composites, estimated intrinsic connectivity with connectome embeddings, and evaluated predictive performance at both Baseline and the 2 Year Follow-Up. For completeness, we benchmarked embeddings against observed and model-implied connectivity in supplementary analyses. Broadly, we found that predictive accuracy was weak at baseline but improved substantially at the two-year follow-up, with differences emerging across substance use facets.

At baseline, predictive models yielded modest effect sizes, typically accounting for less than 1% of the variance in outcomes, consistent with prior literature. By contrast, predictive accuracy was substantially higher for Wave 2 outcomes, with some models explaining over 10% of the variance. Descriptively, intent was the most predictable facet of substance use at baseline, whereas access and family-developmental history were more accurately predicted in 2 Year Follow Up data.

Supplementary analyses showed that node embedding dimensions performed comparably to traditional connectivity metrics, with no consistent advantage for combining feature sets. These findings mark the first connectome-based predictive modeling of substance use to leverage multi-wave ABCD rs-fMRI data, offering new insights into how early brain connectivity forecasts substance use vulnerabilities over time.

### 4.2 Substance Use Intent Was the Most Predictable Facet at Baseline, but Access and Family History Emerged as More Predictive at Follow-Up

At Baseline, the substance use intent facet—which captures both desire to use substances as well as actual use—was the most accurately predicted by rs-fMRI connectomic data. *Post-hoc* analyses predicting strictly substance use (omitting variables about intent to use or curiosity about use) at the 2 Year Follow-Up yielded similarly modest predictive performance, indicating that isolating realized use did not improve prediction relative to broader substance-related composites. This pattern suggests that neural phenotypes may align more closely with latent substance-related tendencies than with infrequent and context-dependent behavioral expression in early adolescence.

In contrast, models performed worst when predicting the access facet at this wave. On the one hand, one could argue this pattern was somewhat unexpected given longstanding research emphasizing the central role of contextual factors—such as peer norms and parental monitoring—in shaping substance use behaviors (Fujimoto & Valente, 2012; Hussong, 2002; Lilja et al., 2003; Steen, 2010; Toumbourou et al., 2007), coupled with work from social neuroscience showing that social dynamics, broadly construed, and their correlates are embedded in network connectivity (Hyon et al., 2020, 2022; Rudolph et al., 2018). One possible explanation is that many of the variables comprising the access composite reflect dynamic, situationally dependent factors—such as peer group behavior or household rules—that may lack stable neurobiological correlates at the onset of adolescence or fluctuate too rapidly to be captured reliably by traditional questionnaire measures. On the other hand, one could argue that it is not unexpected to see weak prediction of access at Baseline, because contextual influences on substance use—such as availability, peer access, or parental restrictions—often become more salient only later in adolescence. At ages 9–10, many youths are still under relatively strong parental oversight, have limited autonomy, and may not yet have substantial exposure to peers who use substances. Under such conditions, the brain may have little to “encode” about access in a way that is stable or predictive. Moreover, one could contend that even when these social dynamics do become influential, it may be somewhat far-fetched to expect them to leave robust, readily detectable signatures in intrinsic connectivity measured with rs-fMRI. With that said, however, this initial weakness in predicting access did not persist, as access became among the most predictable facets at the 2 Year Follow Up.

At the 2 Year Follow-Up, the baseline pattern reversed: both access and family-developmental history were predicted more accurately than intent, with access showing especially strong gains in model performance. In the case of family-developmental history, which was only modestly predicted at baseline, the improved prediction at the 2 Year Follow-Up may reflect the accumulating influence of familial and intergenerational risk factors over time. Although one might expect such biologically grounded influences to manifest more strongly in early brain function, our findings suggest that their effects may unfold more gradually—perhaps due to variability in timing, intensity, or cumulative exposure. By contrast, access—which was poorly predicted at baseline—became among the most predictable outcomes at the 2 Year Follow-Up, aligning with developmental shifts in autonomy and peer exposure that make contextual opportunities for use more salient. This pattern highlights the need to consider both developmental timing and latent growth trajectories when interpreting the connectomic signatures of risk.

It is important to note, however, that the composites were not identical across waves: some variables available at baseline were not collected at the 2 Year Follow-Up due to their inappropriateness for older ages (e.g., substance gating items used in the intent composite at Baseline), leading to slight differences in composite composition.

While this reflects touches on our efforts to maintain the developmental appropriateness of the composites at each time point, it falls in line with a broader, field-wide methodological challenge for longitudinal work: should measures be kept identical across time points to maximize comparability, or titrated to developmental appropriateness (Telzer et al., 2018)? In the present case, the 2 Year Follow-Up composites reflect the latter approach, as we could have carried over scores from items at Baseline but chose not to. This issue should be kept in mind when interpreting differences in predictive performance between waves (we also discuss this issue more in-depth later in the limitations section). Notably the access composite, which showed stark changes in predictive accuracy between Baseline and Follow-Up, comprised the same items at both timepoints. Similarly, trait impulsivity was poorly predicted at both time points, despite continuity in its constituent items. Both sets of findings suggest that continuities or differences in predictive accuracy are not necessarily solely accounted for by changing the composition of composite variables.

Finally, the poor predictability of trait impulsivity—a theoretically important transdiagnostic risk factor for substance use—warrants consideration. One possibility is that the abbreviated UPPS-P used here primarily captures stable, trait-like variance that may not be strongly reflected in intrinsic connectivity patterns at this developmental stage. Alternatively, impulsivity may be better characterized as a state-dependent or context-sensitive construct, whose neural correlates emerge more clearly during task engagement rather than at rest. Measurement limitations may also play a role, as brief self-report scales may insufficiently capture the heterogeneity of impulsivity-related processes relevant for prediction (but see <u>Scheve et al. (2024)</u> as an ABCD-based example to the contrary). Finally, it is possible that neural signatures of impulsivity become more pronounced later in adolescence, suggesting a mismatch between developmental timing of the phenotype and the baseline brain measures used here.

### 4.3 The Default Mode, Somatomotor, and Cingulo-Opercular networks drove predictive accuracy at Follow-Up

At the 2 Year Follow-Up, predictive features were no longer diffusely distributed across the connectome but instead clustered disproportionately within three large-scale systems: the Default Mode, Somatomotor, and Cingulo-Opercular networks. This concentration suggests that as individuals progress into mid-adolescence, the “division of labor” among brain systems relevant to substance use vulnerabilities becomes more focused, with these networks carrying a larger share of predictive signal across outcomes. Importantly, this pattern contrasts with baseline, where weak model performance was accompanied by a diffuse and less interpretable spread of features. Together, these findings indicate that developmental changes are reflected not only in model accuracy but also in the specific network architectures that support prediction.

A closer look at the three networks that dominated predictive features at Year 2 Follow-Up suggests meaningful links to developmental processes relevant for substance use. The default mode network has long been implicated in self-referential thought, social cognition, and affective processing; its prominence here may reflect the increasing role of socioemotional factors, motivational or reward-related drives, and peer-related dynamics in adolescence, which are routinely implicated in substance use initiation (Caouette & Feldstein Ewing, 2017). The cingulo-opercular Network, which supports executive control and salience detection (Gratton et al., 2022), may become more predictive as adolescents’ regulatory capacities are recruited in contexts involving risk, reward, and peer influence. Heightened contributions from this system could reflect the need to detect and manage conflict between long-term goals and immediate opportunities for substance use. The somatomotor network is a less expected contributor (Uddin et al., 2019), but its emergence could signal developmental coupling between sensorimotor processes and substance-related behavior, such as the embodied aspects of substance curiosity or broader integration of motivational and action systems. Although these interpretations are necessarily speculative given the study design, the convergence of these networks underscores that prediction at mid-adolescence reflects the coordinated involvement of systems spanning socioemotional, control, and embodied domains.

### 4.4 Connectome Embeddings Show Promising Applications in Connectome-Based Predictive Modeling

Predictive models trained on node embedding dimensions and model-implied connectivity performed comparably to those based on observed connectivity or combined feature sets. Although these comparisons were conducted as ancillary analyses, the findings are still informative: they suggest node embeddings preserve key topological features of the connectome that are relevant for phenotypic prediction. This comparability both underscores the potential value of connectome embedding approaches and highlights the enduring utility of traditional connectivity measures in predictive modeling. While the similarity in performance may appear trivial at first glance, it carries important implications—namely, that embeddings can be leveraged in future studies without sacrificing predictive fidelity, while also offering advantages in terms of scalability, dimensionality reduction, and potential interpretability.

Our results indicate that node embeddings capture the core predictive signal. In this sense embeddings provide a parsimonious representation of connectivity — not because the models used fewer total predictors (our feature selection procedure always yielded the top 1,000 univariate features in the discovery sample), but because they achieved comparable accuracy without the need for additional feature sets. The lack of performance gains when embeddings were combined with observed or model-implied connectivity suggests that embeddings already capture the core predictive signal. It is in this way that we mean node embeddings represent a parsimonious feature space — they hold their own without the need for additional metrics. Of course, if one *were* interested in building a predictive model with > 1,000 features, then node embeddings provide a more parsimonious framework, in a more traditional sense of the term, for modeling because they reduce the overall number of features (10,560 embedding dimensions overall compared to 61,776 unique connectivity matrix edges). Importantly, equivalent predictive performance should not be interpreted as redundancy, but rather as evidence that explicitly encoding both local and global network structure does not directly confer an obvious performance advantage for this specific predictive task, while still offering representational and analytical benefits beyond raw performance.

Added parsimony of either kind not only promises more computationally and statistically efficient model training, but could also promote conceptual interpretability. Embeddings flexibly capture information about individual brain regions and their implicit ties with other regions, meaning that their inclusion in predictive modeling provides utility for compactly uncovering information about both individual parcels and their network connections. Using node embedding gives an analyst the flexibility to focus their interpretation on individual parcels or networks, or compute embedding-implied connectivity and examine the networks that share the highest implied connectivity with the parcels or network identified in the course of predictive modeling.

In our view, this offers unique opportunities for integrative and longitudinal analyses. For example, embeddings derived from functional connectivity models can be compared to those obtained from structural connectivity (Levakov et al., 2021), enabling direct comparisons across imaging modalities (e.g., identifying parcels from a predictive model trained on functional data, then comparing the functional node embedding for the parcels to an embedding from a structural network). Embeddings could also be considered with multivoxel patterns extracted from the brain regions they represent (Guassi Moreira & Silvers, 2025), providing a richer characterization of neural activity.

Finally, embeddings lend themselves to longitudinal investigations, allowing researchers to track how nodes evolve within the embedding space over time or across developmental stages. Such analyses could offer a conceptually tractable framework for understanding how brain networks adapt with age, experience, and behavioral outcomes. Together, these advantages position node embeddings as a powerful tool for advancing connectome-based modeling, both in terms of substance use prediction and broader developmental neuroscience questions.

### 4.5 Limitations

This study has several limitations that represent noteworthy avenues for future investigation. One limitation of this study lies in our cross-validation scheme. While the use of a single train-test split is efficient and ensures demographic matching across splits, it does carry the risk that the specific split chosen could influence the results, potentially yielding numbers that are not fully representative. However, random splits often result in demographic imbalances that could skew predictive accuracy whereas our demographic matching likely mitigates some of this variability. While a solution could involve generating many demographically matched random splits, this process would be computationally intensive and logistically challenging.

Another limitation relates to the fact that the ABCD dataset is not exhaustibly representative of youth around the world and across time. Future studies could leverage other larger datasets, such as the Philadelphia Neurodevelopmental Cohort or ENGIMA studies (Satterthwaite et al., 2014; Thompson et al., 2020). Relatedly, because substance use behavior evolves as policies, economic circumstances, and other similar factors change, it is important to note that our results may not best describe other cohorts.

A further limitation concerns the construction of the substance-use composites across waves. Several variables that were available at Baseline were not collected at the 2 Year Follow-Up, meaning that the composites are not perfectly matched across time. This issue is hardly unique to our study: developmental cognitive neuroscience more broadly has grappled with the tension between ensuring continuity of measurement and adapting measures for developmental appropriateness (Telzer et al., 2018). On one hand, strict continuity offers the cleanest basis for longitudinal comparison but risks asking questions or administering tasks that may not be age-appropriate or informative as youth mature. On the other hand, titrating measures to developmental stage can enhance sensitivity to the most relevant processes at a given age but comes at the cost of strict cross-wave comparability. The field has yet to converge on best practices given the lack of an obviously “correct” solution to this dilemma. In the present study, we prioritized developmental appropriateness within the constraints of the ABCD dataset. As a result, our findings involving the intent and family-developmental history facets should be interpreted with the understanding that differences in predictive performance across waves may partly reflect differences in measurement as much as changes in underlying neurobehavioral processes.

One last consideration involves our approach to controlling for covariates. Some prior approaches residualize outcomes with respect to their covariates before model training, which to our knowledge could inflate predictive accuracy. Residualization removes variance in the outcome that is shared with covariates but not with the predictors, thereby potentially creating an “easier” prediction problem than would be faced in practice. Relatedly, residualization changes the interpretation of the prediction target: instead of modeling a clinically or behaviorally meaningful phenotype, models are trained to predict a statistically adjusted residual that may not map cleanly onto real-world substance-use behaviors. This shift in the interpretation also affects conclusions about magnitude interpretations: Predicting, say, 8% of a residualized outcome means that less variance is necessarily accounted for when predicting a non-residualized outcome. For these reasons, we included covariates directly in the model with our selected features.

## Acknowledgments

Data used in the preparation of this article were obtained from the Adolescent Brain Cognitive Development (ABCD) Study (abcdstudy.org), held in the NIMH Data Archive (NDA). This is a multisite, longitudinal study designed to recruit more than 10,000 children age 9-10 and follow them over 10 years into early adulthood. The ABCD Study® is supported by the National Institutes of Health and additional federal partners under award numbers U01DA041022, U01DA041028, U01DA041048, U01DA041089, U01DA041106, U01DA041117, U01DA041120, U01DA041134, U01DA041148, U01DA041156, U01DA041174, U24DA041123, and U24DA041147. A full list of supporters is available at https://abcdstudy.org/about/federal-partners/. A listing of participating sites and a complete listing of the study investigators can be found at https://abcdstudy.org/study-sites/. ABCD consortium investigators designed and implemented the study and/or provided data but did not necessarily participate in analysis or writing of this report. This manuscript reflects the views of the authors and may not reflect the opinions or views of the NIH or ABCD consortium investigators. We are also grateful to the NIH-funded Scientific Training in Addiction Research Techniques (START) program for mentorship resources.

## Supplement

### S1. Connectome Embedding Model Details

The node2vec algorithm is based on the seminal word2vec algorithm (Mikolov et al., 2013), originally developed to help estimate vectorized representations of naturalistic speech. While this model was initially used to predict words in a sequence of text from their context, or predicting the context from a target word (Goldberg & Levy, 2014), there is no intrinsic requirement that constrains the algorithm to naturalistic speech and it can be applied to estimate vector embeddings for nodes in a graph object.

The algorithm learns two weight matrices – one that defines the embedding space and contains the vector representations for each node (*W*), and a second one that transforms the embedding layer to the output layer (*W’*). Stochastic gradient descent is used to update these two matrices by minimizing the negative log loss function. Here we used a sliding window of *s* = 3 nodes, meaning that the context nodes were comprised of the *s* nodes that appeared before the target node *and* the *s* nodes that appeared after it. We simulated *o* = 800 random walk sequences, each of length *l* = 20. We specified 30 latent dimensions such that an embedding for a given node contained 30 values.

The random sequences were parameterized such that we applied constants to control the behavior of the walks in order to balance local- vs global-bias. Briefly, communication within functional brain connectivity networks is thought to be driven by local connections between nodes (in a geodesic manner, not necessarily a spatial one). Importantly, the node2vec algorithm assumes information about a node’s local neighborhood, which means the simulated random walks needed to reflect this in order to promote biological and computational compatibility. Consequently, two parameters in the random walk simulation were set to ensure locally-biased sequences. A *p* parameter sets the return probability (walker immediately returns to the node in the next step) and a *q* parameter sets the in-out probability (walker visit a non-connected node in the next step). These constants are divided by the sum of edges depending on the node for the current step in the walk (*p* if the next candidate node is the same as the previous node; *q* if the next candidate node is fully disconnected). Thus, we set *p* = 0.1 and *q* = 1.6 to promote locally-biased networks. These parameters are consistent with prior work using these methods (Rosenthal et al., 2018).

This model fitting procedure was fit to each subject’s functional connectivity data. A random subset of 50 participants had their functional connectivity matrices averaged and used to fit a single connectome embedding model. The embedding space for this particular model was used as a reference space by which all subject-level embedding spaces were subsequently aligned (Levakov et al., 2021). Whereas the embedding similarity of a given node pair is comparable between embedding spaces, the absolute value of the embeddings is not. To ensure comparability between spaces, individual subject embeddings must be aligned to a reference embedding space. This is achieved by multiplying *W’* of the target reference space with the embedding space for the desired participant (*W*).

Next, as a sanity check, we wanted to ensure the embedding model fit properly. We did this by estimating the relationship between observed connectivity and model-implied connectivity that is derived by taking the similarity between two parcel’s embedding vectors. Practically, this means that we took the correlation between each participant’s observed connectivity (correlation matrix comprised of all possible pairwise correlations among the parcel dense time series) and the model implied connectivity obtained by taking the cosine similarity among vector embeddings. This step helped gauge the degree to which the connectome embedding model performs as an accurate representation of the connectivity. Results are shown in Supplementary Figure 1, indicating that the connectome embedding models accurately capture overall rs-fMRI connectivity across participants.

### S2. Variable Details for the Substance Use Composites

The <u>intent</u> composite variable was created in the following manner. First, using data from the baseline timeline follow-back assessment, we created a summed composite of the items indicating whether youths had *heard* about a particular substance (tlfb_ + alc, tob, mj, mj_synth, bitta, caff, inhalant, rx_misuse, list_yes_no). We applied the same procedure to the variables asking youths whether they had *tried* a particular substance (tlfb_alc_sip, tlfb_tob_puff, tlfb_*_use, tlfb_*_use_type *) as well as the variables tapping regular use of each substance (tlfb_*_reg). These three sum score composites (sub_use_heard, sub_use_tried, sub_use_reg) were concatenated with additional variables that tapped alcohol, nicotine, and caffeine use (path_acl_youth[1-8], su_isip_1_calc, su_isip_1b_2_calc, first_nicotine_1, first_nicotine_3, first_mj_1b, su_caff_ss_sum_calc). The final intent composite was created by averaging scores of each individual variable in this concatenated variable set.

The <u>access</u> composite variable was created by computing the mean of the following variables for each participant. peer_deviance_1_4bbe5d, peer_deviance_2_dd1457, peer_deviance_3_e1ec2e, peer_deviance_4_b6c588, peer_deviance_5_bffa44, peer_deviance_6_69562e, peer_deviance_7_beb683, peer_deviance_8_35702e, peer_deviance_9_6dd4ef, parent_rules_q1, parent_rules_q1a, parent_rules_q[2-9], su_risk_p_[1-9].

Finally, the <u>family-developmental history</u> (fdhx) composite was created in the same manner using the following variables. asr_q06_p, asr_q90_p, asr_q124_p, asr_q126_p, scrn_hr_smoke, fam_history_5_yes_no, famhx_4_p, devhx_8_tobacco, devhx_8_alcohol, devhx_8_marijuana, devhx_9_tobacco, devhx_9_alcohol, devhx_9_marijuana.

All items are presented in table form in Supplementary Tables 1A-C; the tables indicate item availability by wave.

---

**Supplementary Figure 1.**
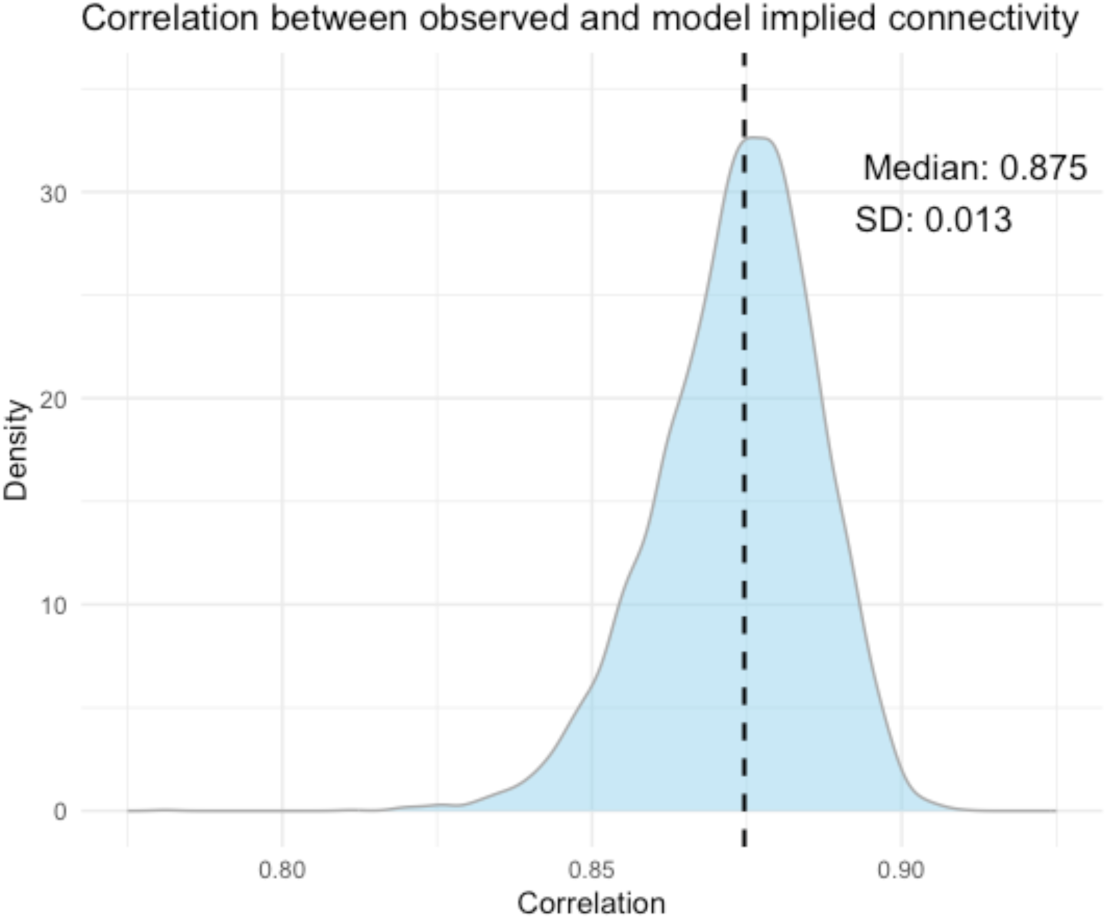
Distribution of observed and model implied connectivity correlations.

**Supplementary Figure 2.**
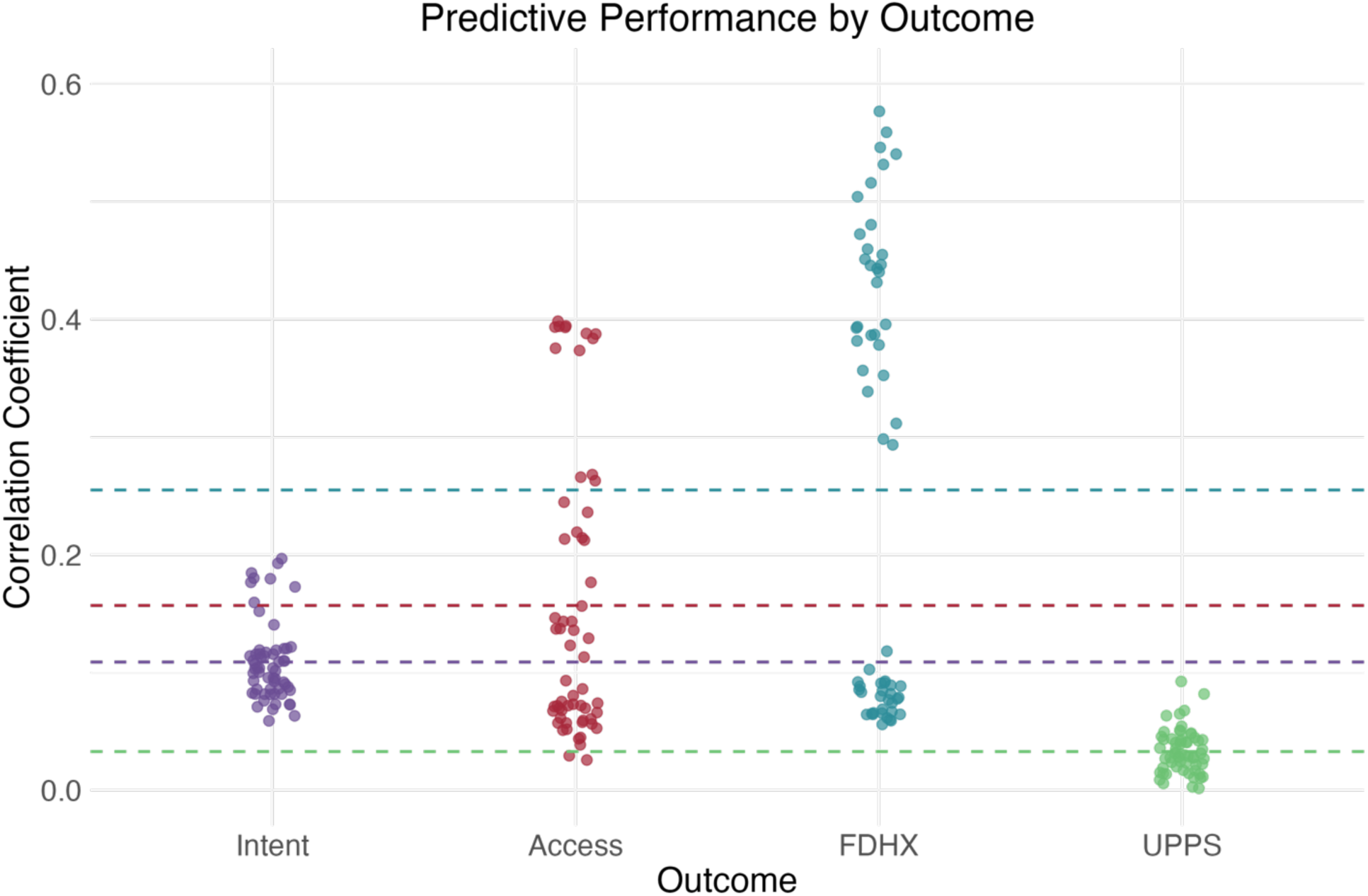
Predictive performances split by outcome, inclusive of all feature sets

**Supplementary Figure 3.**
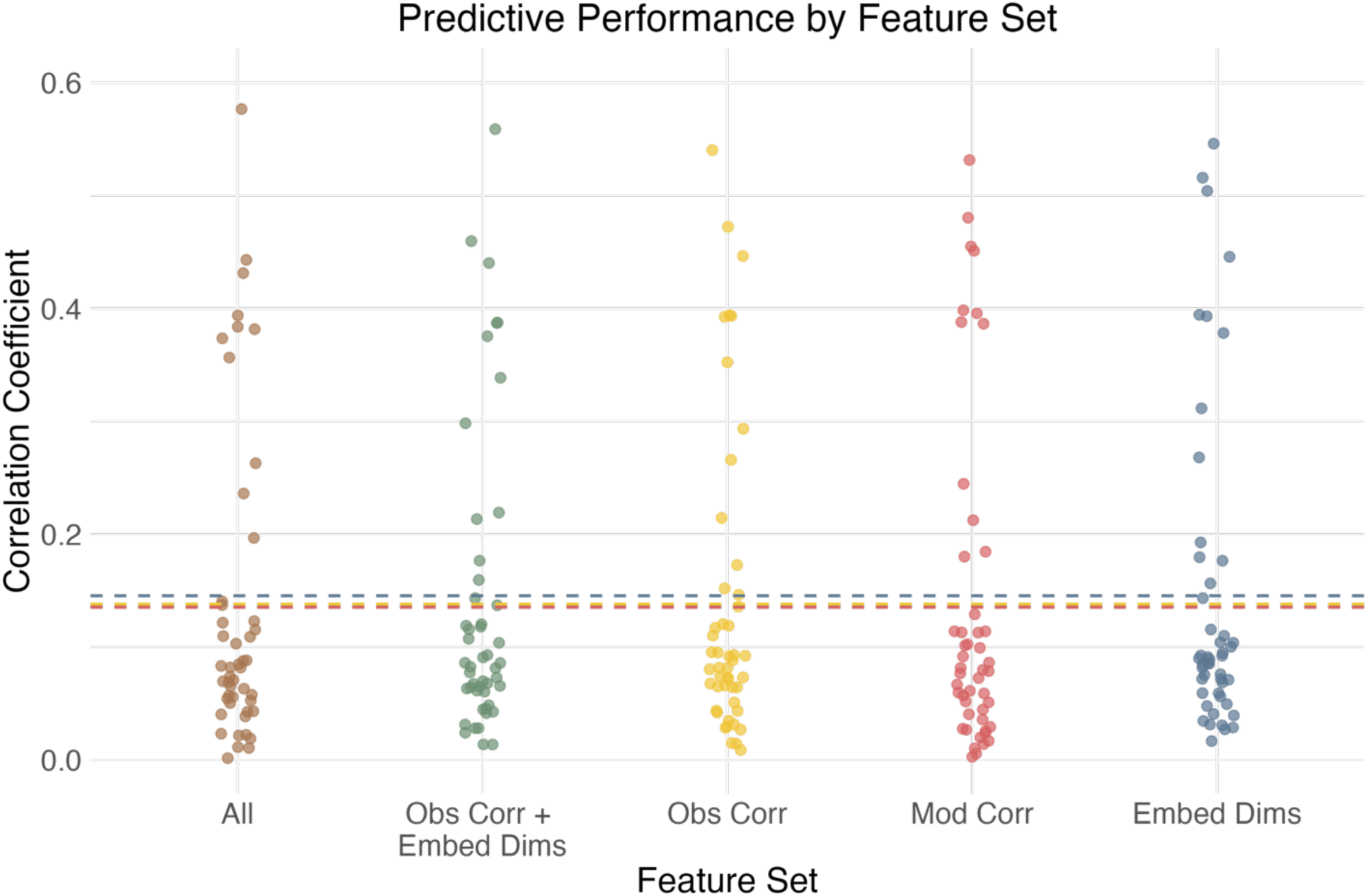
Predictive performances split by all feature sets

**Supplementary Figure 4.**
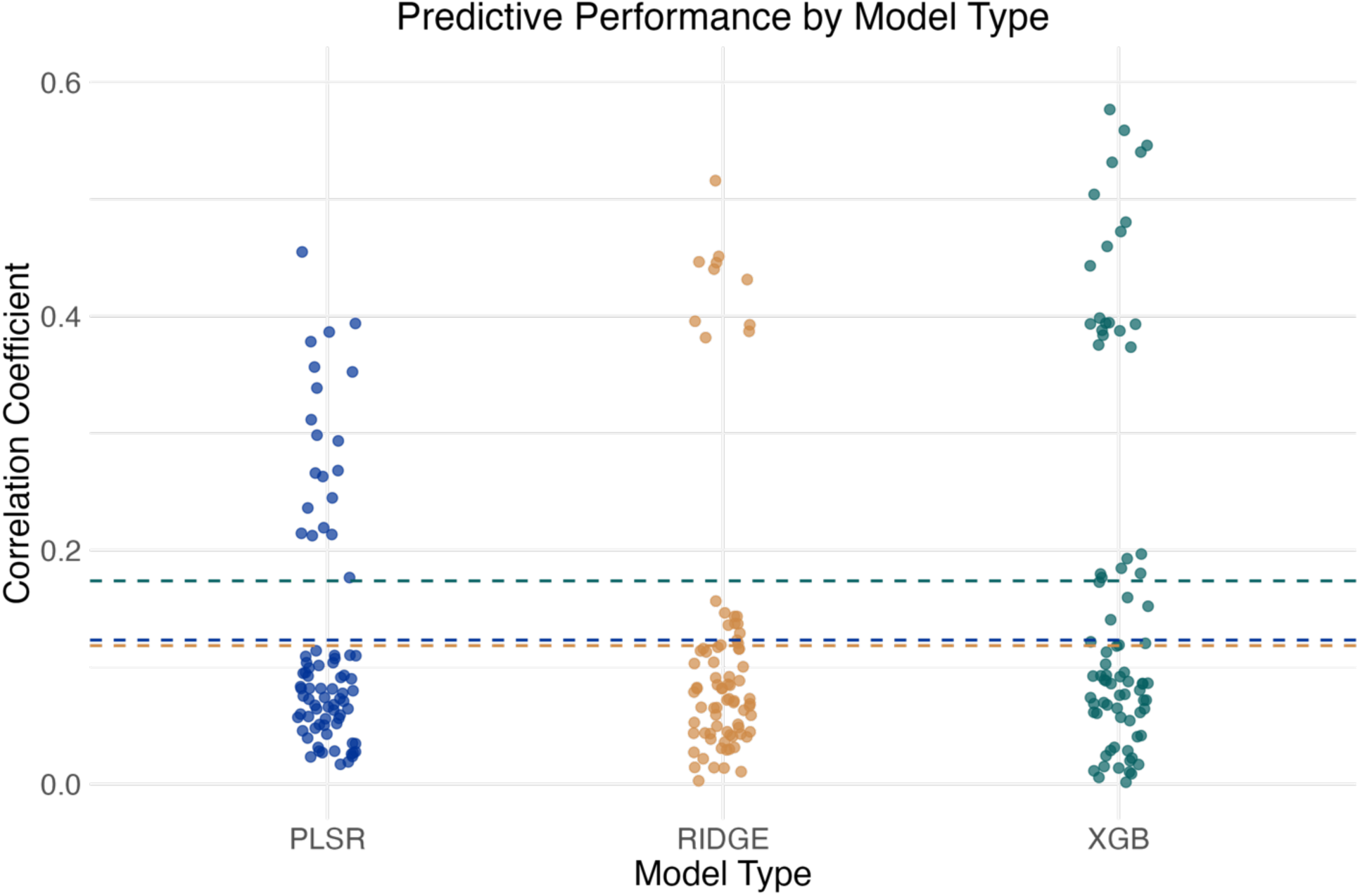
Predictive performances by model types, inclusive of all feature sets

**Supplementary Figure 5.**
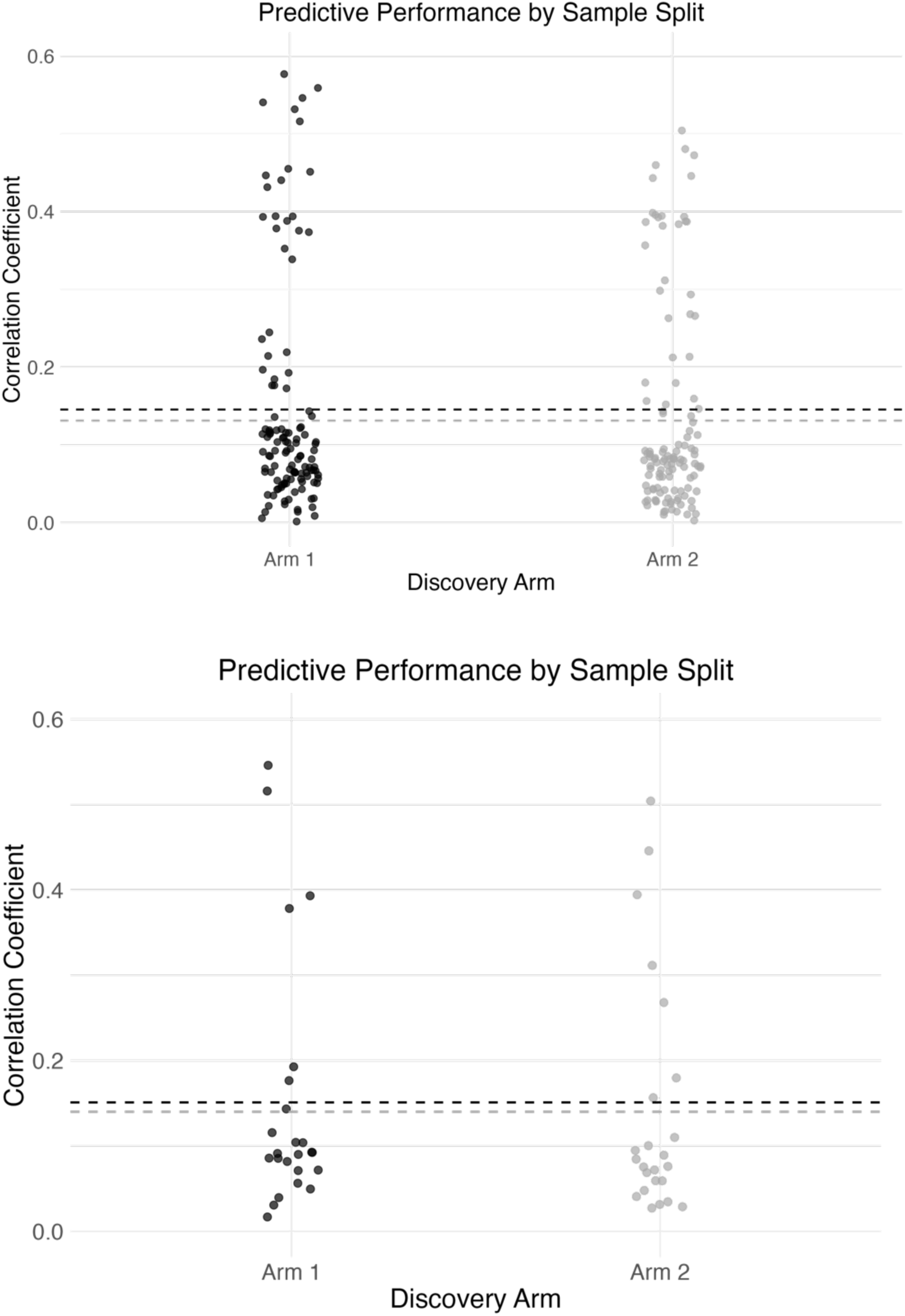
Predictive performances plotted by which arm served as the confirmation dataset (top: inclusive of all feature sets, bottom: model specifications with node embedding feature set only)

**Supplementary Table 1A.**
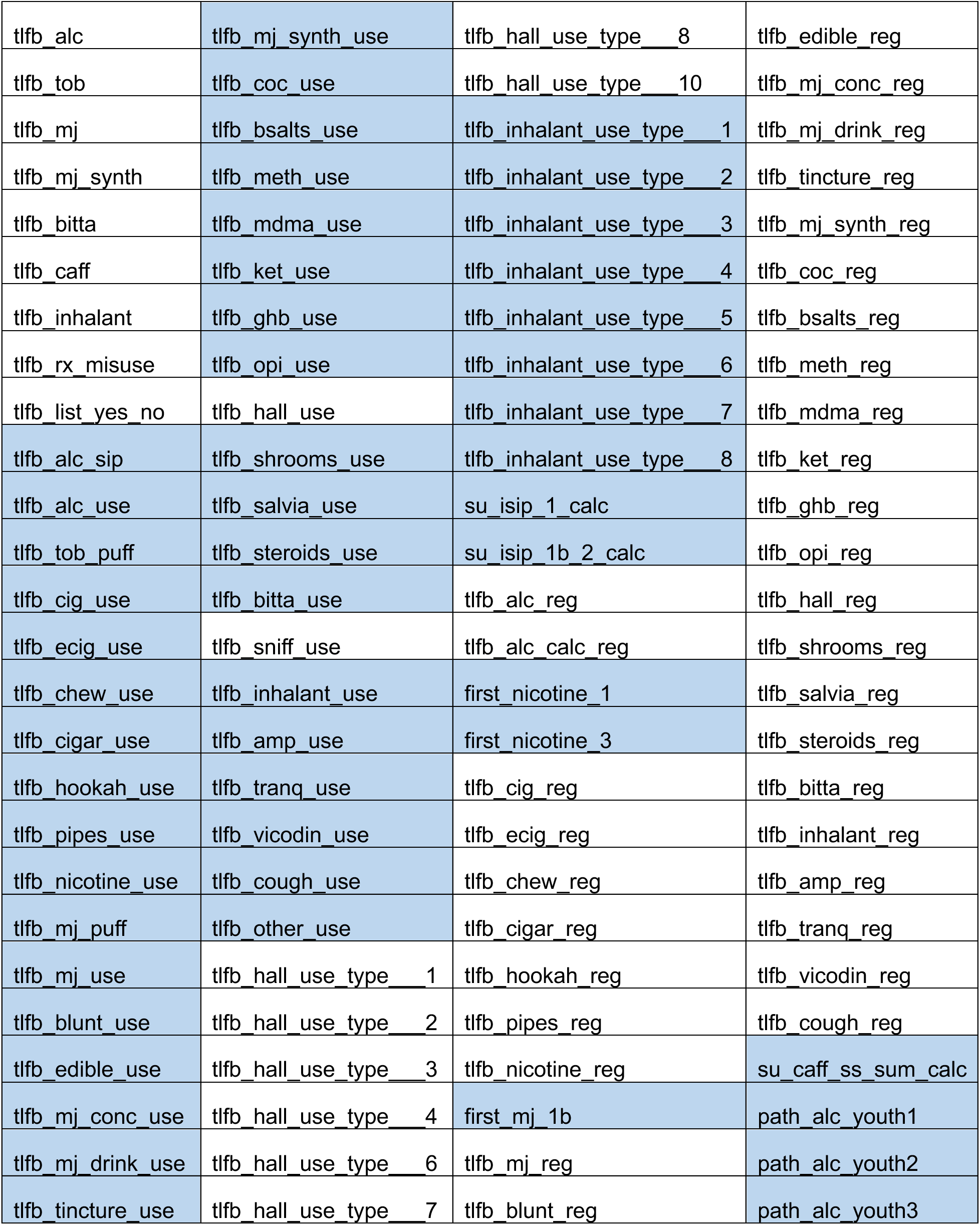
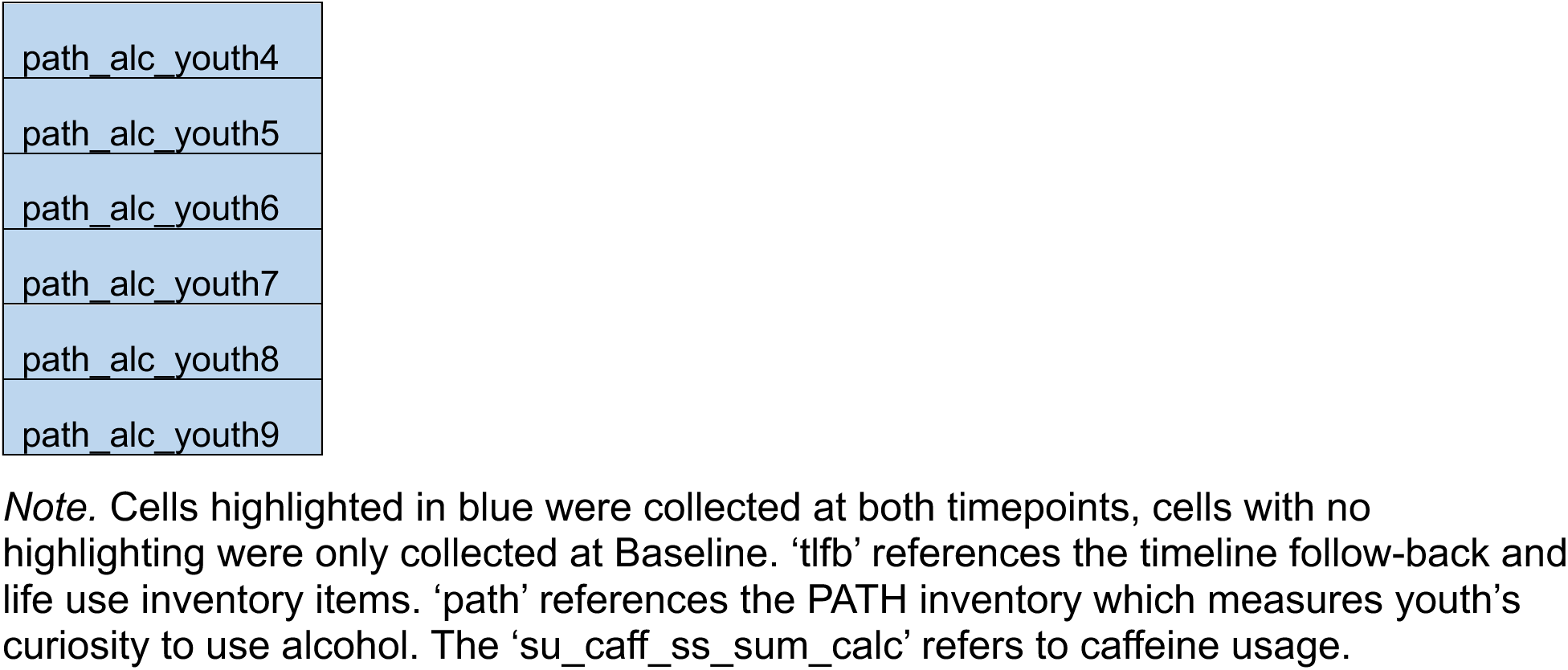
Full set of *intent* variables used in the composite

|  |  |  |  |
| --- | --- | --- | --- |
| tlfb_alc | tlfb_mj_synth_use | tlfb_hall_use_type_8 | tlfb_edible_reg |
| tlfb_tob | tlfb_coc_use | tlfb_hall_use_type_10 | tlfb_mj_conc_reg |
| tlfb_mj | tlfb_bsalts_use | tlfb_inhalant_use_type_1 | tlfb_mj_drink_reg |
| tlfb_mj_synth | tlfb_meth_use | tlfb_inhalant_use_type_2 | tlfb_tincture_reg |
| tlfb_bitta | tlfb_mdma_use | tlfb_inhalant_use_type_3 | tlfb_mj_synth_reg |
| tlfb_caff | tlfb_ket_use | tlfb_inhalant_use_type_4 | tlfb_coc_reg |
| tlfb_inhalant | tlfb_ghb_use | tlfb_inhalant_use_type_5 | tlfb_bsalts_reg |
| tlfb_rx_misuse | tlfb_opi_use | tlfb_inhalant_use_type_6 | tlfb_meth_reg |
| tlfb_list_yes_no | tlfb_hall_use | tlfb_inhalant_use_type_7 | tlfb_mdma_reg |
| tlfb_alc_sip | tlfb_shrooms_use | tlfb_inhalant_use_type_8 | tlfb_ket_reg |
| tlfb_alc_use | tlfb_salvia_use | su_isip_1_calc | tlfb_ghb_reg |
| tlfb_tob_puff | tlfb_steroids_use | su_isip_1b_2_calc | tlfb_opi_reg |
| tlfb_cig_use | tlfb_bitta_use | tlfb_alc_reg | tlfb_hall_reg |
| tlfb_ecig_use | tlfb_sniff_use | tlfb_alc_calc_reg | tlfb_shrooms_reg |
| tlfb_chew_use | tlfb_inhalant_use | first_nicotine_1 | tlfb_salvia_reg |
| tlfb_cigar_use | tlfb_amp_use | first_nicotine_3 | tlfb_steroids_reg |
| tlfb_hookah_use | tlfb_tranq_use | tlfb_cig_reg | tlfb_bitta_reg |
| tlfb_pipes_use | tlfb_vicodin_use | tlfb_ecig_reg | tlfb_inhalant_reg |
| tlfb_nicotine_use | tlfb_cough_use | tlfb_chew_reg | tlfb_amp_reg |
| tlfb_mj_puff | tlfb_other_use | tlfb_cigar_reg | tlfb_tranq_reg |
| tlfb_mj_use | tlfb_hall_use_type_1 | tlfb_hookah_reg | tlfb_vicodin_reg |
| tlfb_blunt_use | tlfb_hall_use_type_2 | tlfb_pipes_reg | tlfb_cough_reg |
| tlfb_edible_use | tlfb_hall_use_type_3 | tlfb_nicotine_reg | su_caff_ss_sum_calc |
| tlfb_mj_conc_use | tlfb_hall_use_type_4 | first_mj_1b | path_alc_youth1 |
| tlfb_mj_drink_use | tlfb_hall_use_type_6 | tlfb_mj_reg | path_alc_youth2 |
| tlfb_tincture_use | tlfb_hall_use_type_7 | tlfb_blunt_reg | path_alc_youth3 |

|  |
| --- |
| path_alc_youth4 |
| path_alc_youth5 |
| path_alc_youth6 |
| path_alc_youth7 |
| path_alc_youth8 |
| path_alc_youth9 |

*Note.* Cells highlighted in blue were collected at both timepoints, cells with no highlighting were only collected at Baseline. ‘tlfb’ references the timeline follow-back and life use inventory items. ‘path’ references the PATH inventory which measures youth’s curiosity to use alcohol. The ‘su\_caff\_ss\_sum\_calc’ refers to caffeine usage.

**Supplementary Table 1B.** Full set of *Access* variables used in the composite

|  |  |  |  |
| --- | --- | --- | --- |
| peer_deviance_1_4bbe5d | peer_deviance_8_35702e | parent_rules_q5 | su_risk_p_3 |
| peer_deviance_2_dd1457 | peer_deviance_9_6dd4ef | parent_rules_q6 | su_risk_p_4 |
| peer_deviance_3_e1ec2e | parent_rules_q1 | parent_rules_q7 | su_risk_p_5 |
| peer_deviance_4_b6c588 | parent_rules_q1a | parent_rules_q8 | su_risk_p_6 |
| peer_deviance_5_bffa44 | parent_rules_q2 | parent_rules_q9 | su_risk_p_7 |
| peer_deviance_6_69562e | parent_rules_q3 | su_risk_p_1 | su_risk_p_8 |
| peer_deviance_7_beb683 | parent_rules_q4 | su_risk_p_2 | su_risk_p_9 |

*Note.* Cells highlighted in blue were collected at both timepoints, cells with no highlighting were only collected at Baseline.

**Supplementary Table 1C.** Full set of *FDHX* variables used in the composite

|  |  |  |
| --- | --- | --- |
| asr_q06_p | fam_history_5_yes_no | devhx_9_tobacco |
| asr_q90_p | famhx_4_p | devhx_9_alcohol |
| asr_q124_p | devhx_8_tobacco | devhx_9_marijuana |
| asr_q126_p | devhx_8_alcohol |  |
| scrn_hr_smoke | devhx_8_marijuana |  |

*Note.* Cells highlighted in blue were collected at both timepoints, cells with no highlighting were only collected at Baseline.

**Supplementary Table 2.**
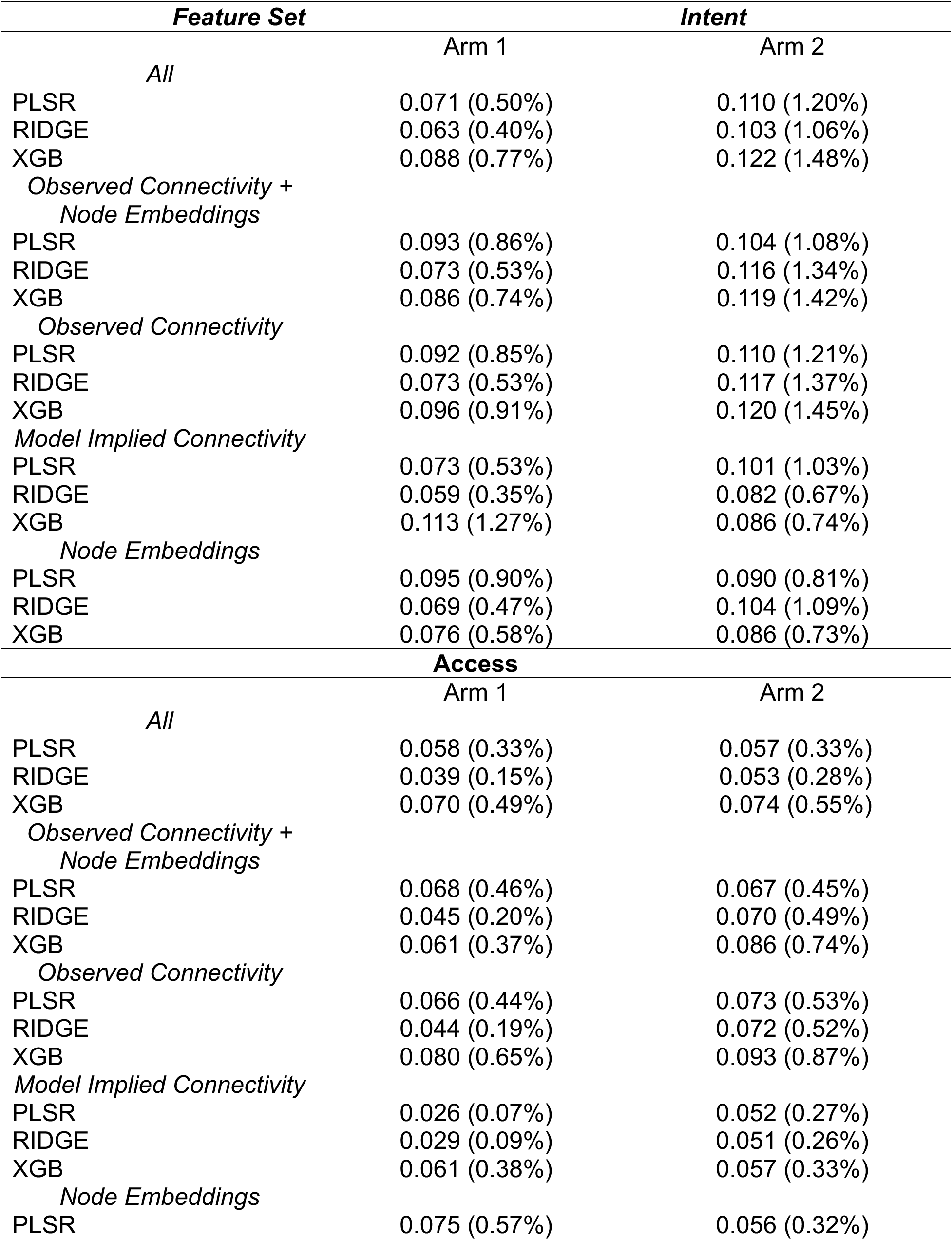
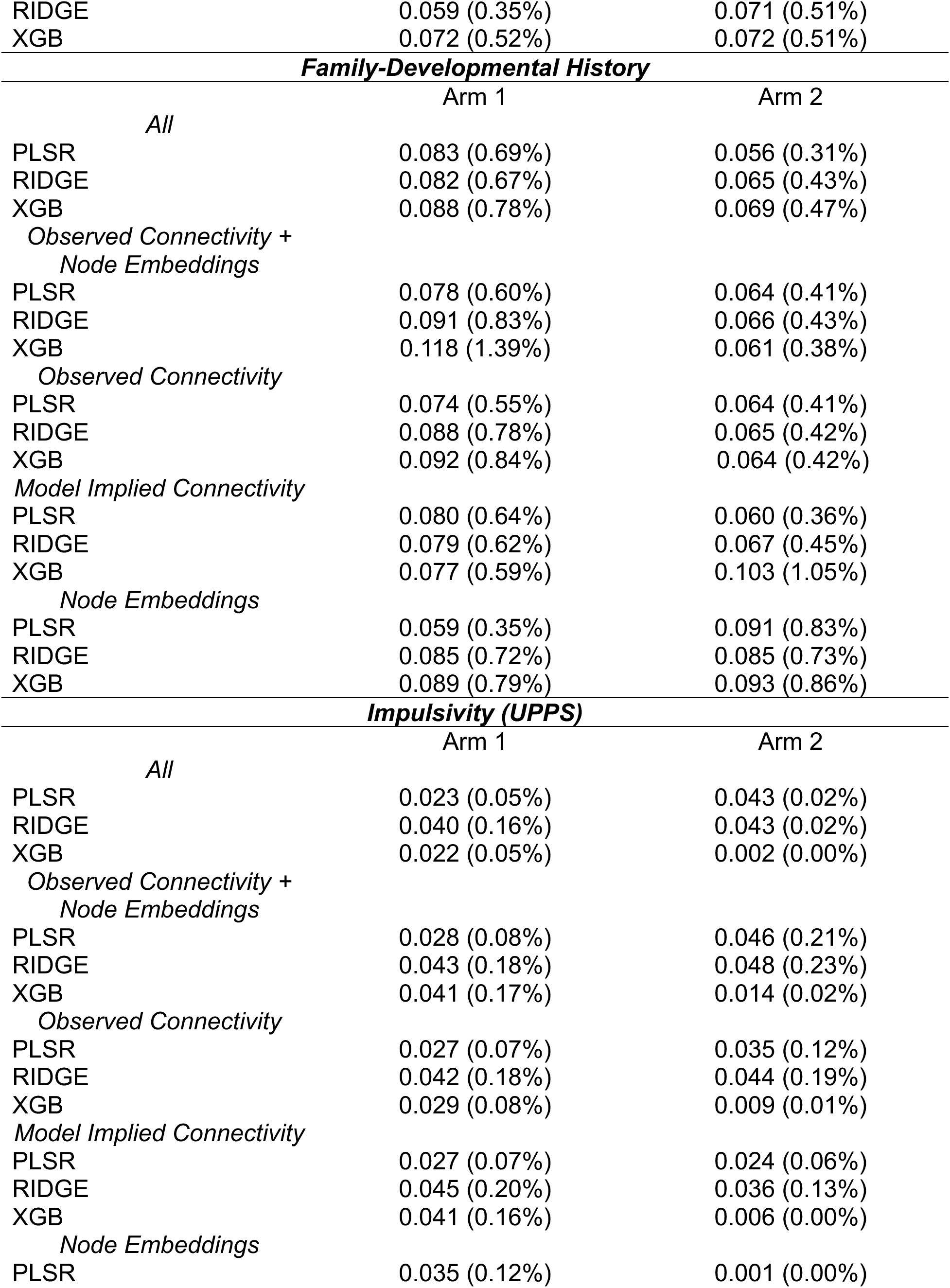
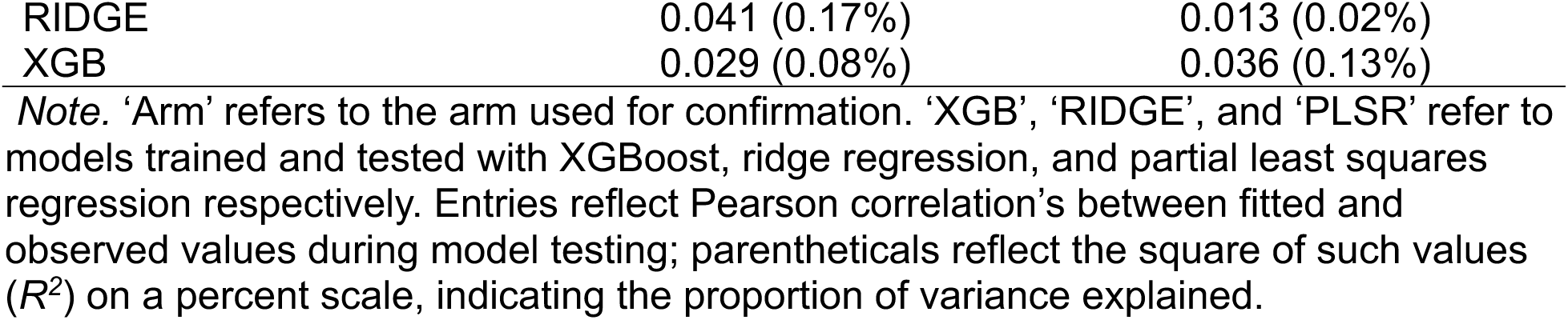
Model performances predicting substance use composites at Baseline broken down by all possible feature combinations

**Supplementary Table 3.** Model performances predicting substance use composites at the 2 Year Follow-Up broken down by all possible feature combinations

| **Intent** |  |  |
| --- | --- | --- |
|  | Arm 1 | Arm 2 |
| *All* |  |  |
| PLSR | 0.082 (0.67%) | 0.109 (1.19%) |
| RIDGE | 0.085 (0.72%) | 0.116 (1.33%) |
| XGB | 0.141 (1.97%) | 0.197 (3.87%) |
| *Observed Connectivity +*<br>*Node Embeddings* |  |  |
| PLSR | 0.081 (0.66%) | 0.107 (1.15%) |
| RIDGE | 0.083 (0.68%) | 0.120 (1.45%) |
| XGB | 0.159 (2.54%) | 0.177 (3.12%) |
| *Observed Connectivity* |  |  |
| PLSR | 0.082 (0.67%) | 0.095 (0.91%) |
| RIDGE | 0.082 (0.67%) | 0.119 (1.41%) |
| XGB | 0.152 (2.31%) | 0.173 (2.98%) |
| *Model Implied Connectivity* |  |  |
| PLSR | 0.099 (0.99%) | 0.114 (1.3%) |
| RIDGE | 0.092 (0.84%) | 0.114 (1.3%) |
| XGB | 0.180 (3.25%) | 0.185 (3.4%) |
| *Node Embeddings* |  |  |
| PLSR | 0.110 (1.21%) | 0.104 (1.08%) |
| RIDGE | 0.100 (1.01%) | 0.116 (1.34%) |
| XGB | 0.180 (3.23%) | 0.193 (3.72%) |
| **Access** |  |  |
|  | Arm 1 | Arm 2 |
| *All* |  |  |
| PLSR | 0.263 (6.91%) | 0.236 (5.57%) |
| RIDGE | 0.137 (1.88%) | 0.123 (1.51%) |
| XGB | 0.384 (14.73%) | 0.374 (13.95%) |
| *Observed Connectivity +*<br>*Node Embeddings* |  |  |
| PLSR | 0.213 (4.55%) | 0.219 (4.80%) |
| RIDGE | 0.143 (2.05%) | 0.137 (1.88%) |
| XGB | 0.388 (15.02%) | 0.375 (14.10%) |
| *Observed Connectivity* |  |  |
| PLSR | 0.266 (7.07%) | 0.214 (4.60%) |
| RIDGE | 0.146 (2.14%) | 0.136 (1.85%) |
| XGB | 0.393 (15.48%) | 0.394 (15.53%) |
| *Model Implied Connectivity* |  |  |
| PLSR | 0.212 (4.51%) | 0.245 (5.99%) |
| RIDGE | 0.129 (1.67%) | 0.113 (1.28%) |
| XGB | 0.398 (15.87%) | 0.388 (15.05%) |
| *Node Embeddings* |  |  |
| PLSR | 0.268 (7.19%) | 0.177 (3.12%) |
| RIDGE | 0.156 (2.45%) | 0.143 (2.06%) |
| XGB | 0.394 (15.56%) | 0.393 (15.46%) |
| **Family-Developmental History** |  |  |
|  | Arm 1 | Arm 2 |
| *All* |  |  |
| PLSR | 0.357 (12.71%) | 0.394 (15.51%) |
| RIDGE | 0.382 (14.57%) | 0.431 (18.61%) |
| XGB | 0.443 (19.63%) | 0.577 (33.26%) |
| *Observed Connectivity +*<br>*Node Embeddings* |  |  |
| PLSR | 0.298 (8.90%) | 0.339 (11.47%) |
| RIDGE | 0.387 (14.99%) | 0.440 (19.39%) |
| XGB | 0.460 (21.13%) | 0.559 (31.24%) |
| *Observed Connectivity* |  |  |
| PLSR | 0.293 (8.61%) | 0.352 (12.42%) |
| RIDGE | 0.393 (15.42%) | 0.446 (19.93%) |
| XGB | 0.472 (22.32%) | 0.540 (29.20%) |
| *Model Implied Connectivity* |  |  |
| PLSR | 0.387 (14.94%) | 0.455 (20.70%) |
| RIDGE | 0.396 (15.66%) | 0.451 (20.35%) |
| XGB | 0.480 (23.08%) | 0.532 (28.26%) |
| *Node Embeddings* |  |  |
| PLSR | 0.312 (9.71%) | 0.378 (14.31%) |
| RIDGE | 0.446 (19.87%) | 0.516 (26.62%) |
| XGB | 0.504 (25.42%) | 0.546 (29.82%) |
| **Impulsivity (UPPS)** |  |  |
|  | Arm 1 | Arm 2 |
| *All* |  |  |
| PLSR | 0.019 (0.04%) | 0.050 (0.25%) |
| RIDGE | 0.011 (0.01%) | 0.022 (0.05%) |
| XGB | 0.011 (0.01%) | 0.054 (0.29%) |
| *Observed Connectivity +*<br>*Node Embeddings* |  |  |
| PLSR | 0.028 (0.08%) | 0.063 (0.40%) |
| RIDGE | 0.014 (0.02%) | 0.031 (0.10%) |
| XGB | 0.024 (0.06%) | 0.065 (0.42%) |
| *Observed Connectivity* |  |  |
| PLSR | 0.031 (0.10%) | 0.051 (0.26%) |
| RIDGE | 0.014 (0.02%) | 0.030 (0.09%) |
| XGB | 0.015 (0.02%) | 0.068 (0.46%) |
| *Model Implied Connectivity* |  |  |
| PLSR | 0.017 (0.03%) | 0.028 (0.08%) |
| RIDGE | 0.003 (0.00%) | 0.014 (0.02%) |
| XGB | 0.010 (0.01%) | 0.020 (0.04%) |
| *Node Embeddings* |  |  |
| PLSR | 0.048 (0.23%) | 0.082 (0.67%) |
| RIDGE | 0.027 (0.07%) | 0.031 (0.09%) |
| XGB | 0.031 (0.10%) | 0.092 (0.85%) |

**Supplementary Table 4.** Model performances predicting an aggregate of all substance use variables at the 2 Year Follow-Up, broken down by all possible feature combinations

|  | *Use* |  |
| --- | --- | --- |
|  | Arm 1 | Arm 2 |
| *All* |  |  |
| PLSR | 0.026 (0.07%) | 0.001 (0.00%) |
| RIDGE | 0.030 (0.09%) | 0.013 (0.02%) |
| XGB | 0.042 (0.18%) | 0.036 (0.13%) |
| *Observed Connectivity +*<br>*Node Embeddings* |  |  |
| PLSR | 0.018 (0.03%) | 0.012 (0.01%) |
| RIDGE | 0.025 (0.06%) | 0.016 (0.03%) |
| XGB | 0.025 (0.06%) | 0.048 (0.23%) |
| *Observed Connectivity* |  |  |
| PLSR | 0.018 (0.03%) | 0.012 (0.01%) |
| RIDGE | 0.023 (0.05%) | 0.018 (0.03%) |
| XGB | 0.028 (0.08%) | 0.026 (0.07%) |
| *Model Implied Connectivity* |  |  |
| PLSR | 0.021 (0.05%) | 0.010 (0.01%) |
| RIDGE | 0.023 (0.05%) | 0.023 (0.05%) |
| XGB | 0.052 (0.27%) | 0.017 (0.03%) |
| *Node Embeddings* |  |  |
| PLSR | 0.062 (0.38%) | 0.045 (0.20%) |
| RIDGE | 0.043 (0.18%) | 0.024 (0.06%) |
| XGB | 0.034 (0.11%) | 0.020 (0.04%) |

*Note.* ‘Arm’ refers to the arm used for confirmation. ‘XGB’, ‘RIDGE’, and ‘PLSR’ refer to models trained and tested with XGBoost, ridge regression, and partial least squares regression respectively. Entries reflect Pearson correlation’s between fitted and observed values during model testing; parenteticals reflect the square of such values ( $R^2$ ) on a percent scale, indicating the proportion of variance explained.

**Supplementary Table 5.** Model performances predicting an alternate calculation of the family developmental history composite at the 2 Year Follow-Up, broken down by all possible feature combinations

| ***Family-Developmental History, Alternate*** |  |  |
| --- | --- | --- |
|  | Arm 1 | Arm 2 |
| *All* |  |  |
| PLSR | 0.447 (19.97%) | 0.510 (25.99%) |
| RIDGE | 0.446 (19.87%) | 0.486 (23.58%) |
| XGB | 0.579 (33.57%) | 0.613 (37.56%) |
| *Observed Connectivity +*<br>*Node Embeddings* |  |  |
| PLSR | 0.408 (16.66%) | 0.462 (21.34%) |
| RIDGE | 0.456 (20.79%) | 0.495 (24.45%) |
| XGB | 0.578 (33.36%) | 0.631 (39.86%) |
| *Observed Connectivity* |  |  |
| PLSR | 0.427 (18.23%) | 0.475 (22.59%) |
| RIDGE | 0.460 (21.18%) | 0.499 (24.89%) |
| XGB | 0.576 (33.20%) | 0.618 (38.23%) |
| *Model Implied Connectivity* |  |  |
| PLSR | 0.465 (21.63%) | 0.542 (29.37%) |
| RIDGE | 0.462 (21.37%) | 0.505 (25.49%) |
| XGB | 0.589 (34.75%) | 0.665 (44.22%) |
| *Node Embeddings* |  |  |
| PLSR | 0.362 (13.11%) | 0.445 (19.79%) |
| RIDGE | 0.514 (26.39%) | 0.557 (31.00%) |
| XGB | 0.610 (37.16%) | 0.637 (40.60%) |

*Note.* ‘Arm’ refers to the arm used for confirmation. ‘XGB’, ‘RIDGE’, and ‘PLSR’ refer to models trained and tested with XGBoost, ridge regression, and partial least squares regression respectively. Entries reflect Pearson correlation’s between fitted and observed values during model testing; parentheticals reflect the square of such values ( $R^2$ ) on a percent scale, indicating the proportion of variance explained. The alternate calculation of the composite includes family history items that were measured at baseline and were not included in the main formulation of the composite for the 2 Year Follow-Up.

## Notes

### Competing Interest Statement

The authors have declared no competing interest.

### Summary of Updates

This version of the manuscript has been revised in response to several concerns following peer review, including requests for better clarity, re-focusing the discussion away from speculative clinical implications, and the inclusion of additional follow-up analyses.

